# Opposing Nodal and Wnt signalling activities govern the emergence of the mammalian body plan

**DOI:** 10.1101/2025.01.11.632562

**Authors:** André Dias, Pau Pascual-Mas, Gaëlle Robertson, Gabriel Torregrosa-Cortés, Suzan Stelloo, Pablo Casaní-Galdón, Stephen Babin, Yuliia Romaniuk, Alexandre Mayran, Alexandra E. Wehmeyer, Jordi Garcia-Ojalvo, Harold M. McNamara, Michiel Vermeulen, Sebastian J. Arnold, Alfonso Martinez Arias

**Affiliations:** Department of Medicine and Life Sciences, Universitat Pompeu Fabra, Spain; Barcelona Institute for Global Health, Spain; EMBL Barcelona, Spain; Department of Molecular Biology, Faculty of Science, Radboud Institute for Molecular Life Sciences, Oncode Institute, Radboud University Nijmegen, the Netherlands; School of Life Sciences, Ecole Polytechnique Fédérale de Lausanne (EPFL), Switzerland; Institute of Experimental and Clinical Pharmacology and Toxicology, Faculty of Medicine, University of Freiburg, Germany; Lewis-Sigler Institute, Princeton University, USA; Wu Tsai Institute and Department of Molecular, Cellular, and Developmental Biology, Yale University, USA; Division of Molecular Genetics, Netherlands Cancer Institute, the Netherlands; Signalling Research Centers BIOSS and CIBSS, University of Freiburg, Germany; ICREA, Spain

## Abstract

Nodal and Wnt signalling play an important role in the emergence of the mammalian body plan, primarily by orchestrating gastrulation. While the literature suggests they cooperate to build the primitive streak, their individual contributions remain poorly understood. Using gastruloids, we found that Wnt/β-catenin drives a genetic program characteristic of the late primitive streak, promoting the development of posterior body structures in a time and dose-dependent manner. Conversely, Nodal activates a distinct transcriptional module resembling the early streak. By engineering gastruloids with varying levels of Nodal signalling, we demonstrate that a decreasing temporal gradient of Nodal activity is critical for establishing the anterior body, with higher Nodal levels producing more anterior structures in a concentration-dependent manner. Our findings suggest that Nodal and Wnt act antagonistically, initiating distinct developmental modules within the primitive streak. This antagonism is likely the core mechanism driving the early body plan in mammals.

## Introduction

The Primitive Streak (PS) is a transient structure that emerges at the midline of the posterior epiblast in mammals during gastrulation and gives rise to the primordia of the main organs of the embryo, except for the brain^1,2^. Fate mapping and lineage tracing studies have shown that the early PS is configured by cells located initially in the proximal-posterior part of the epiblast, that undergo an epithelial to mesenchymal transition (EMT) and give rise, in an ordered sequence, to extraembryonic mesoderm, anterior mesoderm (cranial and cardiac) and endoderm^3–9^. As gastrulation proceeds, some of these cells persist in the anterior/distal-most part of the streak while a new pool from the anterior half of the epiblast emerges at the posterior compartment forming what is referred to as the late posterior PS^4,7,10^. In addition, while the early PS contribute to the node/notochord, cells from the late PS will be incorporated into the caudal epiblast and generate post-occipital neural and mesodermal derivatives during axial extension^4,10–13^. Understanding gastrulation requires understanding the regulation of these cell fate assignment events and their interactions.

It has been known for a long time that mutations of the T-box transcription factor *Eomesodermin* (*Eomes*) result in the loss of derivatives of the early PS^14^ with *Brachyury* (*Tbxt*) mutants revealing a complementary phenotype for the late PS streak, with a loss of axial progenitors and post-occipital tissue^12,15^. Recent studies have shown the reason for this: Eomes suppresses the expression of Tbxt targets, and it is the downregulation of *Eomes* at E7.5 that enables the activation of the late PS program^16^. In mammals, gastrulation is under the control of Nodal and Wnt/β- catenin signalling activity in the posterior part of the epiblast^9,17–21^. While it is known that these signals interact with *Eomes* and *Tbxt* during the laying down of the body plan, the nature of this relationship and its impact on the fate map that emerges during gastrulation remain open questions.

Here we use ‘gastruloids’, a stem cell-based model of mammalian gastrulation and early axial elongation^22–25^, to explore the role of Wnt and Nodal signalling in mammalian gastrulation. When treated with CHIR (CHIR99021), a Gsk3 inhibitor/β-catenin signalling agonist, mouse gastruloids exhibit patterns of gene expression and organisation characteristic of the late PS^24,26,27^. However, these gastruloids exhibit low and variable levels of expression of genes associated with the early PS reflected, in particular, in varying levels of endoderm^27–30^.

Here we show that replacing CHIR for the Nodal surrogate Activin A (ActA) elicits early PS fates with little of the posterior PS-associated ones. This is reflected in the robust expression of *Eomes*, *Foxa2* and *Lhx1*, the absence of significant *Hox* gene expression, and the emergence of anterior mesoderm and pharyngeal endoderm. The observation that Nodal and Wnt/β-catenin drive two alternative genetic programmes in the PS, provides a rationale for the distinct activity dynamics of these signalling in the mammalian gastrula^20,21^ and suggests the existence of two functional developmental modules: a Nodal/*Eomes*, governing anterior fates and a Wnt/*Tbxt*, driving posterior ones. Furthermore, we noticed that while high Nodal and low Wnt levels initiate the anterior mesendodermal program, the transition to low Nodal and high Wnt activates posterior fates. In this transition and interactions, we uncover a potential role for Wnt signalling in pacing the process of gastrulation and the orderly establishment of the mammalian embryonic body plan.

## Results

### Wnt signalling regulates the cell state transition driving the emergence of multipotency in mammals

During the first 48h of the standard mouse gastruloid protocol (**Methods**), cells undergo major transcriptomic changes at a bulk level as they transition from naïve to formative/primed pluripotency. These changes are associated with the downregulation of genes associated with naïve pluripotency (*Zfp42/Rex1*, *Nanog*, *Essrb* and *Tfcp2l1*)^31–34^ and the acquisition of an early epiblast-like signature^4,35^ characterised by the expression of *Fgf5*, *Otx2*, *Pou5f1/Oct4* and *Utf1* (**Fig. 1** and **Extended Data Fig. 1a**). At this stage, gastruloids resemble epiblast-like stem cells (EpiLSCs) and model the pre-streak epiblast at around embryonic day (E) 5.5/E6.0^2,4,36^, displaying significant levels of *Nodal* and limited *Eomes*, *Wnt3* and *Tbxt* expression (**Fig. 1c** and **Extended Data Fig. 1a**). Upon treatment with the standard CHIR concentration (3μM), 72h gastruloids display an upregulation of *Tbxt*, *Cdx2*, *Fst*, *Wnt3a*, *Wnt8a* and *Epha1* (**Fig. 1c**), which are genes normally associated with the emergence of the caudal epiblast at about E8.0 in the embryo^12,37,38^. At the same time, we observed a downregulation of early epiblast-related genes and low expression of *Eomes* (**Fig. 1c**), suggesting that the early PS program is being suppressed and that these gastruloids represent the late PS.

**Fig 1.**
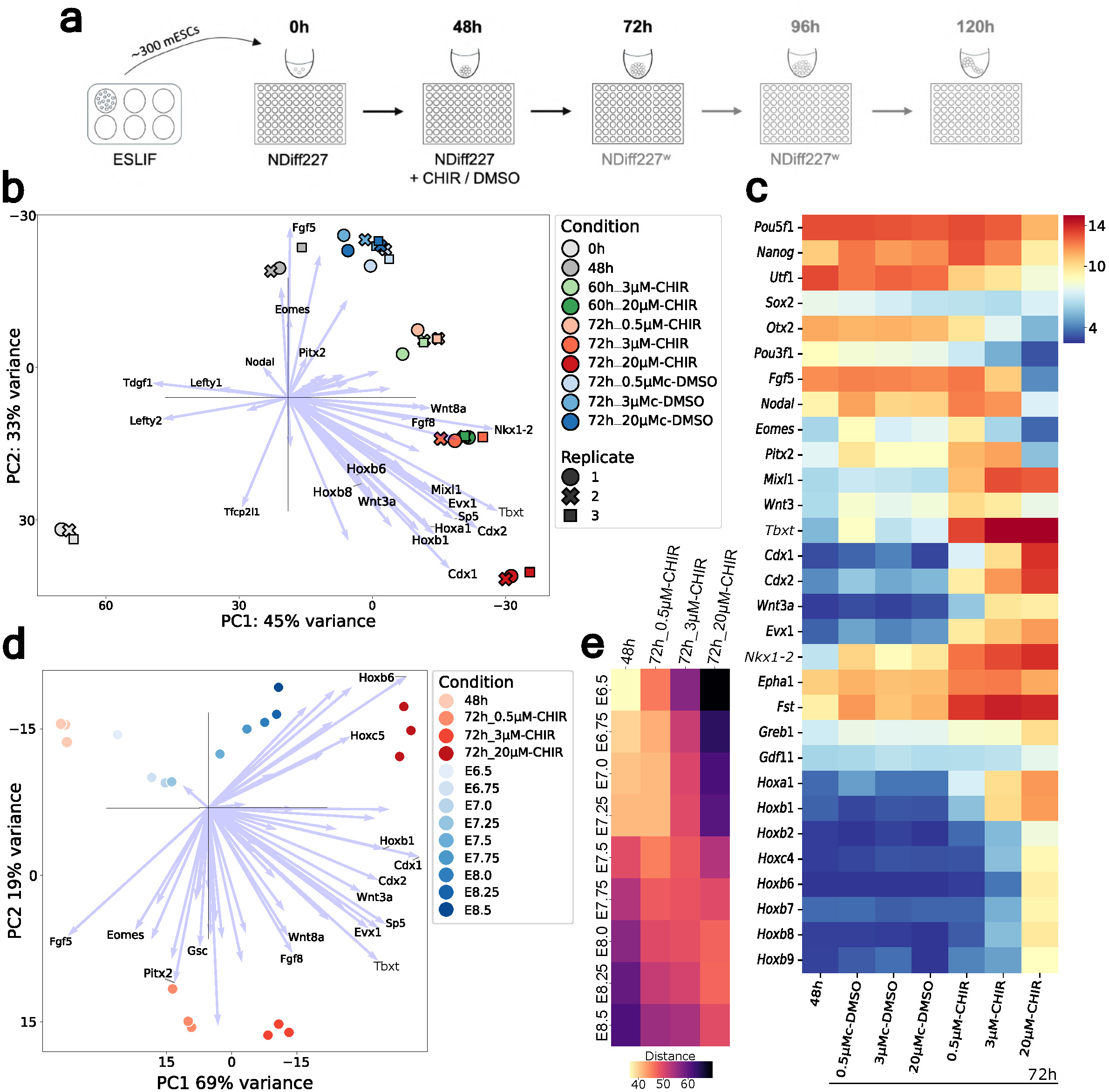
Wnt levels modulate the temporal activation of the late posterior primitive streak. **a**, Schematic diagram of the mouse gastruloid protocol using different CHIR concentrations and their corresponding DMSO controls. The protocol starts with the seeding of approximately 300 mouse embryonic stem cells (mESCs) cultured in ESLIF media (see **Methods** section), as single cells, per well of a 96-well plate in N2B27 media (0h). Following aggregation, CHIR+DMSO (DMSO is vehicle) or just DMSO are introduced at 48h in fresh N2B27. Gastruloids were then collected for bulk RNAseq at various time points (three replicates per condition, from five independent experiments; total gastruloid numbers are indicated in the **Methods** section). **b**, PCA of distinct temporal gastruloid bulk RNA sequencing samples, with some of the top 60 genes derived from the loadings of components 1 and 2 (**Supplementary Data Figure 4a**). Vectors representing genes associated with naive pluripotency (e.g. *Tfcp2l1*) are directed towards the 0h replicates, which cluster far apart from the other conditions. Early epiblast markers such as *Nodal*, *Eomes*, *Fgf5* and *Pitx2*, are represented in loading vectors pointing towards the 48h and DMSO conditions, indicating a change in cell state from 0 to 48h (see also **Extended Data Fig. 1a**). DMSO replicates/conditions cluster together, suggesting that varying DMSO concentrations do not significantly alter the gastruloid transcriptome at 72h. Genes related to the late posterior PS (e.g. *Tbxt*, *Wnt8a* and *Cdx2*) are in loading vectors pointing along the CHIR conditions, which capture its concentration dependence. The 72h gastruloids treated with 3µM CHIR cluster between those treated with 0.5 and 20µM CHIR along this axis. Gastruloids at 60h treated with 3µM CHIR cluster near the 72h gastruloids treated with 0.5µM CHIR, whereas those at 60h that were treated with 20µM CHIR cluster close to 72h gastruloids treated with 3µM CHIR. Together, this highlights the temporal and concentration-dependent effects of Wnt signalling on gastruloids progression along PC1, which represents developmental time. **c**, - Gene expression heatmap (mean variance-stabilized read counts of replicates) showing that the early epiblast-like signature observed at 48h and in DMSO conditions (e.g. *Utf1*, *Fgf5* and *Otx2*) gradually disappears with the increase of CHIR levels. In contrast, expression of genes related to the late PS (e.g. *Tbxt*, *Evx1* and *Fst*) increases with CHIR exposure. This transition is more pronounced in 72h gastruloids treated with 20µM CHIR, which display a strong downregulation of pluripotent and early epiblast/PS markers (e.g. Nanog and *Eomes*, respectively) and a concomitant upregulation of genes related to the caudal epiblast (e.g. *Cdx2*, *Nkx1-2* and *Greb1*). **d,** Pseudo-bulk, time-resolved points of scRNA-seq mouse embryo gastrulation atlas^86^ (blue dots, see **Methods** for the pseudo-bulk construction) projected onto PCA of various CHIR gastruloids. The projection is robust to the analysis parameters (see PCA robustness analysis in **Extended Data Fig. 1b**). Gastruloids at 48h cluster near E6.5 mouse embryos, whereas 72h gastruloids treated with 20µM CHIR cluster towards the opposite end, closer to E8.0-8.5 mouse embryos. 0,5 and 3µM CHIR-treated gastruloids cluster halfway along dimension 1 in a concentration-dependent manner, reflecting the temporal progression of mouse embryos. Among the top 50 gene loadings from components 1 and 2 (**Supplementary Data Figure 4b**), the presence of early epiblast and late/caudal epiblast markers (e.g. *Eomes* and *Cdx2*, respectively) pointing to opposite ends of dimension 1 further validates the temporal developmental changes occurring in the mouse epiblast during gastrulation. **e,** Euclidean distance between the gastruloid and embryo (pseudo-bulk atlas^86^) datasets suggests that the transcriptome of 48h gastruloids closely resembles that of E6.5 mouse embryos, whereas 72h gastruloids more closely resemble later embryonic stages in a concentration-dependent manner. The increase in distance for later time points is likely due to the increase of complexity in mouse cell diversity in comparison to gastruloids.

To better understand the role of Wnt signalling during mammalian gastrulation, we treated mouse gastruloids with different levels of CHIR (**Extended Data Fig. 1c**) and compared their transcriptional signatures through bulk RNA sequencing. Using principal component analysis (PCA), we observed that gastruloids treated with different CHIR concentrations cluster separately along PC1, in a concentration-dependent manner (**Fig. 1b**). Importantly, we noticed that low concentrations of CHIR (0,5μM) resulted in a gene expression profile more akin to the early PS^4,35^ with low levels of *Tbxt*, *Evx1* and *Cdx2*, and high levels of *Eomes*, *Fgf5* and *Utf1*, while a higher concentration of CHIR (20μM) led to a gene signature characteristic of the caudal lateral epiblast (CLE)^12^ with high *Cdx1/2*, *Nkx1-2*, *Wnt3a* and *Greb1*, and low *Nodal*, *Eomes*, *Wnt3* and *Fgf5* (**Fig. 1c**). This resemblance to the caudal epiblast was confirmed through bioinformatic comparison against mouse embryo data (**Fig. 1c** and **Extended Data Fig. 1b**) and can also be observed in the expression of *Hox* genes, like *Hoxb9*, that are prematurely expressed relative to standard gastruloids (**Fig. 1c**). As controls for the variations in CHIR concentrations, we analysed gastruloids treated only with the corresponding levels of the CHIR vehicle (DMSO). Independent of the concentration, these gastruloids clustered together in the PCA and displayed an epiblast-like signature similar to that of 48h gastruloids and EpiLSCs (**Fig. 1** and **Extended Data Fig. 1a**).

These results indicate that Wnt signalling drives the emergence of a late PS/caudal epiblast genetic program in a dose-dependent manner and suggest two possible interpretations. According to the first one, which is in line with standard views of Wnt signalling and gene expression, different levels of Wnt/β-catenin elicit distinct transcriptomic profiles. A second interpretation would suggest that the CLE emerges within a schedule in gene expression whose tempo is modulated by Wnt signalling: the higher the level of Wnt signalling, the faster the progress through the schedule. To distinguish between these possibilities, we treated gastruloids with varying levels of CHIR and collected them at 60h for bulk RNAseq. PCA analysis indicated that, at this time, standard gastruloids cluster closer to 0,5μM CHIR-treated gastruloids at 72h, whereas 20μM CHIR gastruloids cluster near standard gastruloids at 72h (**Fig. 1b**). This observation favours the second interpretation. Further evidence for this is provided by HCR of different epiblast markers showing how different concentrations of CHIR control their expression in mouse gastruloids and how that matches with what happens in the embryo (**Fig. 1** and **Extended Data Fig. 2**).

An observation from the transcriptomic comparison of gastruloids treated with different CHIR levels is the possible relationship between increasing levels of Wnt signalling and early signs of EMT^39^: downregulation of *Cdh1* and increased *Cdh2* and *Snai1* expression (**Extended Data Fig. 3a**). However, this does not correlate with substantial changes in the expression of early mesodermal markers, such as *Tbx6*, *Mesp1* or *Tcf15* (**Extended Data Fig. 3**) which might be expected from a classical cell fate decision model. Given the substantial reduction in pluripotency-associated genes in gastruloids exposed to high Wnt levels (e.g. *Oct4* and *Utf1*) (**Fig. 1c**), our data suggest that Wnt signalling works primarily to control the pace of a state transition leading to multipotency rather than directly acting on fate specification.

Notably, Wnt signalling does not seem to directly downregulate the pluripotent state because, in addition to the maintenance of Oct4 expression, we observed an immediate short-term increase in the levels of Nanog in response to low and standard CHIR treatments (**Fig. 1c** and **2)**. Through 3D segmentation and quantification analysis of gastruloids stained for gastrulation markers, we observed that this increase in the levels of Nanog occurs concomitantly with the gradual upregulation of both Cdx2 and Tbxt (**Fig. 2** and **Extended Data Fig. 4**). Given the co-expression of Nanog and Cdx2, particularly at 72h, and the downregulation of Otx2 (**Fig. 2** and **Extended Data Fig. 4**), this process mediated by the CHIR treatment appears to mimic the maturation of the posterior epiblast of the embryo at the beginning of gastrulation^40–42^. These events precede the emergence of axial progenitors, including the neuromesodermal competent population^13^, which in standard CHIR-treated gastruloids occurs around 96h^24,26,27,43^, providing a rationale for why Wnt signalling is required for gastrulation. In agreement, we noticed that *Wnt3* mutant gastruloids not treated with CHIR do not undergo such epiblast maturation and gradually start differentiating towards a more pro-neural, Sox1^+^ anterior epiblast-like identity (**Fig. 2** and **Extended Data Fig. 4** and 5). These gastruloids diverge from their WT counterparts (WT-DMSO), which undergo symmetry breaking later, around 96h, in a variable and unorganised manner (**Extended Data Fig. 5**). Together, these results explain how Wnt signalling drives the emergence of multipotency by initiating gastrulation and triggering the late PS, and imply that additional factors likely regulate the early PS and the specification of anterior body structures in mammals.

**Fig 2.**
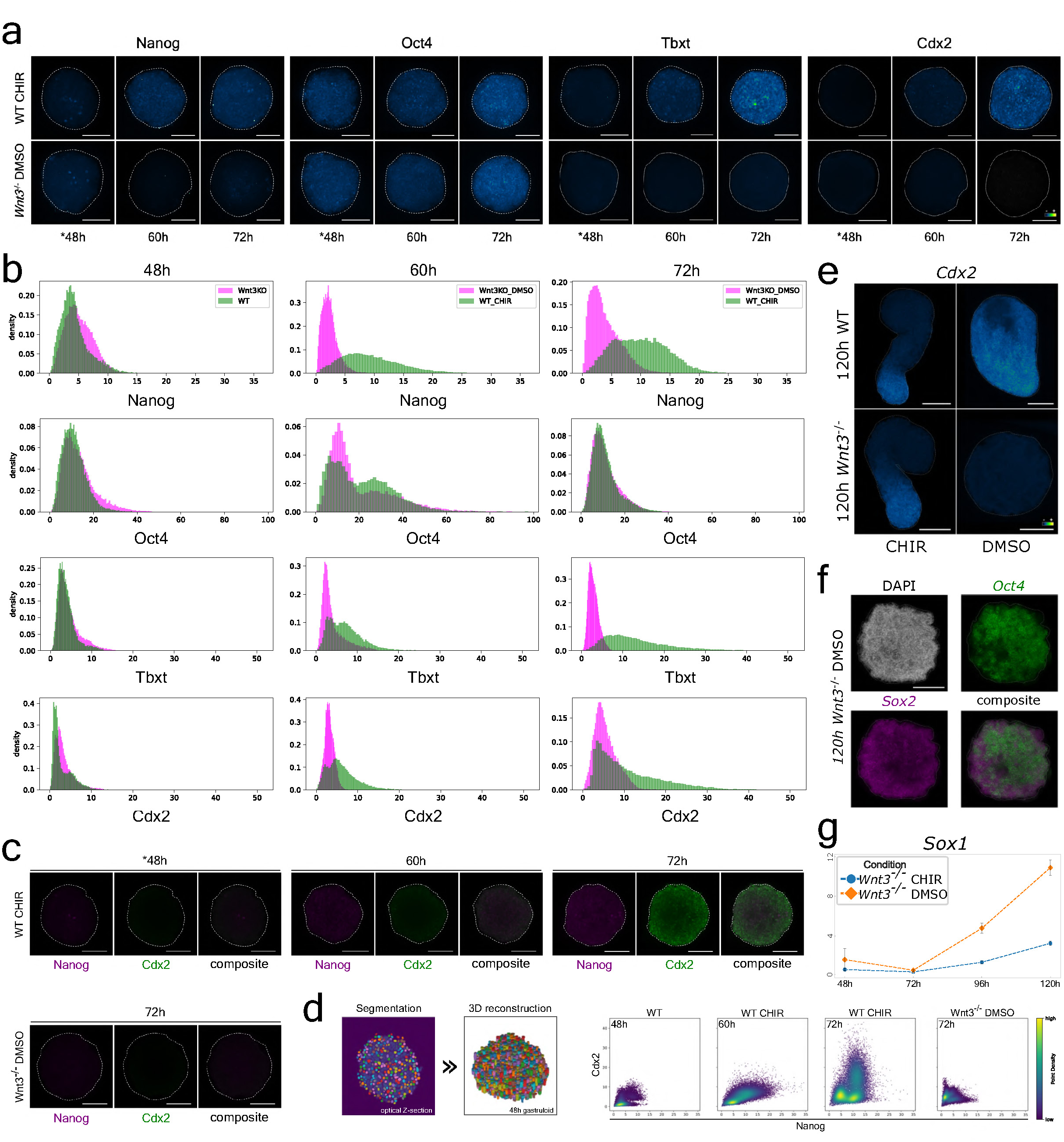
Wnt signalling is required for the acquisition of a posterior epiblast pluripotency-like state in gastruloids. **a**, Maximum intensity projections (MIP) of immunofluorescence stainings (4-5 biological replicates) for Nanog, Oct4, Cdx2 and Tbxt in WT-CHIR and *Wnt3*KO– DMSO treated gastruloids (see also **Extended Data Fig. 4** and 5, and the **Methods** for the experiment details). All treatments were given between 48h and 72h (*48h gastruloids were not yet treated). The expression of these markers is similar across 48h WT and *Wnt3* mutant gastruloids: Oct4 is expressed throughout the gastruloids, Tbxt and Cdx2 are not expressed, and Nanog is only expressed in a handful of cells. In contrast, key differences exist between the two types of gastruloids at 60h and 72h. For instance, Tbxt and Cdx2 expression increases in a temporal manner in WT-CHIR gastruloids but continue to be absent in Wnt3 mutant DMSO treated gastruloids. Nanog expression increases in gastruloids treated with CHIR and is absent/very low in *Wnt3*^-/-^ DMSO gastruloids. Oct4 levels remain consistent between the two conditions. The scale bars are equivalent to 100µm. **b**, Histograms displaying the quantification of antibody signal intensity observed across 3D segmented cells from the WT and *Wnt3* mutant gastruloids shown in (a). There is an increase of Nanog, Cdx2 and Tbxt over time in WT gastruloids treated with CHIR, and their levels are higher than in the mutant gastruloids treated with DMSO. **c**, 20μm optical section Z-stacks showing the expression of Nanog (magenta) and Cdx2 (green) in WT gastruloids treated with CHIR and non-treated *Wnt3* mutant gastruloids. Their expression increases over time in WT gastruloids, and co-localisation (in whitish/greyish) can be detected at the external part of the gastruloids at 60h and in broader domains at 72h. The scale bars are equivalent to 100µm. **d**, Scatter plots highlight the expression of Nanog and Cdx2 in single cells over time in 3D segmented WT and mutant gastruloids, corrected for signal drift along the Z-axis (see **Methods** and **Supplementary Data Fig.1-3**). **e**, HCR for *Cdx2* in 120h WT or *Wnt3*^-/-^ CHIR and DMSO treated gastruloids (treatments between 48 and 72h). Similar to WT-CHIR gastruloids, *Wnt3*KO-DMSO gastruloids display the standard anterior-posterior (A-P) elongation, with *Cdx2* being expressed in the posterior compartment. WT-DMSO gastruloids exhibit a deficient/incomplete elongation process (significant inter-gastruloid variability was observed, with ∼50% displaying an ovoid shape), with *Cdx2* being expressed across the majority of the gastruloid though more enhanced in one of the compartments. In contrast, no expression of *Cdx2* was observed in *Wnt3^-/-^* DMSO gastruloids. They do not elongate and exhibit a round shape at 120h (see also **Extended Data Fig. 5a**). Images (MIP) shown here are a representative selection from 4-5 gastruloids imaged per condition, two independent experiments. The scale bar is equivalent to 200µm. **f**, Characterisation of *Wnt3* mutant gastruloids (also in **Extended Data Fig. 5b**) via HCR for *Oct4* (green) and *Sox2* (in magenta), DAPI is shown in grey. The mutant gastruloids display high levels of expression of these genes at 120h, with extensive co-expression in several compartments (white/greyish colour in composite). A total of 4 gastruloids were imaged, from two independent experiments. The scale bar is equivalent to 200µm. **g**, Plot showing the expression of *Sox1* via qPCR in *Wnt3* mutant gastruloids, treated with either DMSO or CHIR between 48h and 72h (three replicates). *Wnt3* mutant gastruloids treated with DMSO (in orange) display an increase in *Sox1* expression between 72h and 120h that is around 3 times higher than the one noticed in CHIR-treated gastruloids (in blue), characteristic of neural differentiation. Dashed lines between timepoints represent linear interpolations to aid visualisation.

### Gastruloids engineered with Activin A model the early primitive streak

Given the key roles that Nodal signalling plays in gastrulation^18,19,44^, we wondered what would happen if we substituted Wnt signalling for a pulse of Nodal signalling in gastruloids. To do this, we added ActA instead of CHI between 48 and 72h (**Fig. 3a**). An initial comparison of the gene expression patterns in these gastruloids through HCR revealed a robust expression of both *Eomes* and *Tbxt* at 72h, which is consistent with the activation of the early PS program. By 96h, we could detect in the periphery of these gastruloids, the expression of genes related to endoderm (*Foxa2* and *Sox17*) and anterior mesoderm (*Mesp1* and *Tbx6*), which are fates known to arise from the early PS. In addition, we detected node/notochord-like cells (*Noto* expression) in these gastruloids (**Fig. 3f** and **Extended Data Fig. 6**). These patterns of gene expression contrast with those in CHIR-treated gastruloids that only express *Tbxt* at 72h and do not contain signs of notochord or endodermal-like cells at 96h (**Fig. 3f**).

**Fig. 3.**
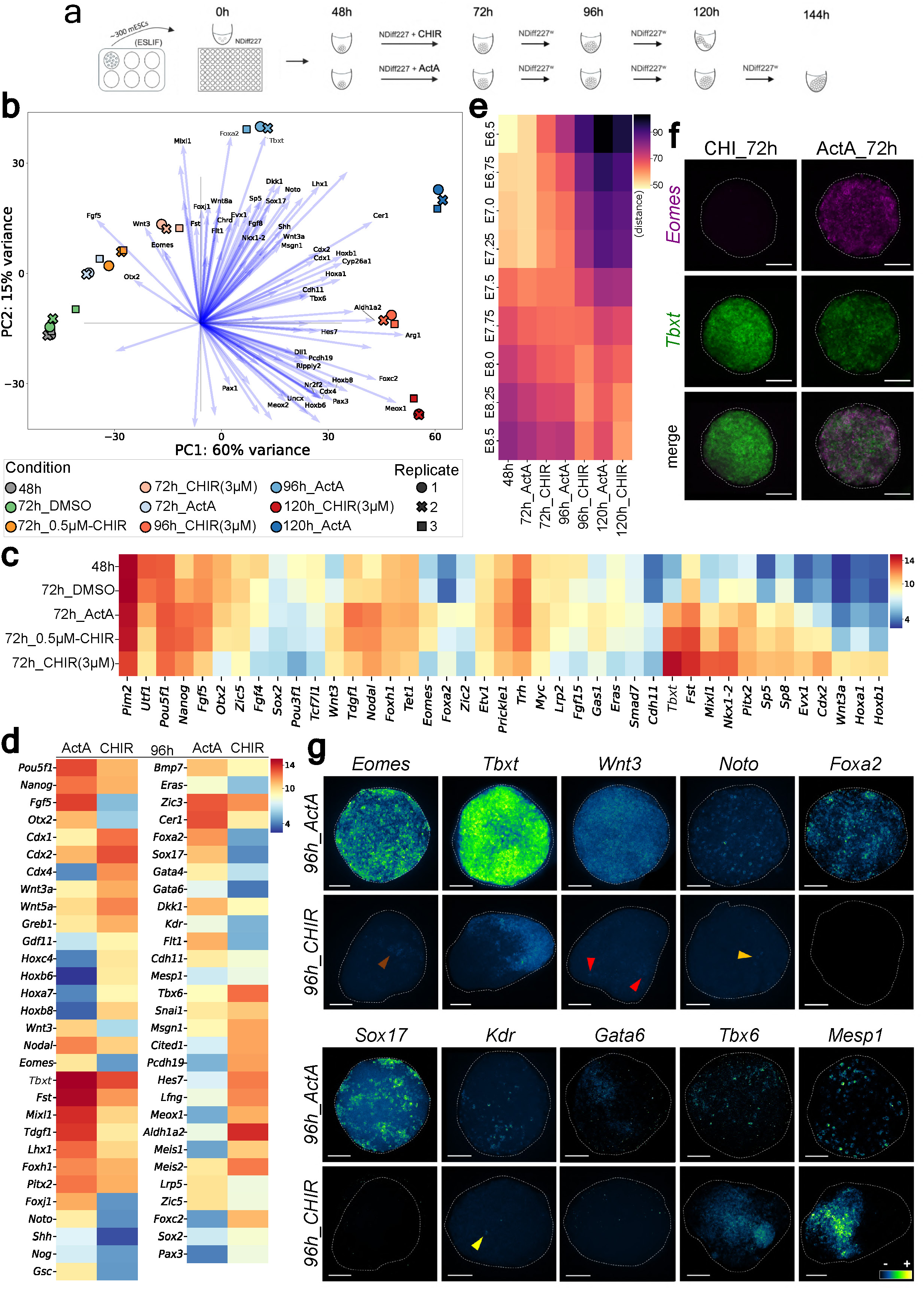
The gastruloid as a stem cell-based embryo model of the primitive streak. **a,** Schematic diagram of the gastruloid engineering process with either CHIR or ActA. Standard CHIR-treated gastruloids exhibit an elongated phenotype at 120h, their endpoint. Conversely, gastruloids made with ActA develop until 144h and remain largely spherical. **b,** PCA of temporal ActA and CHIR-treated gastruloid bulk RNAseq samples (three replicates per condition, from five independent experiments), with some of the top 100 genes derived from the loadings of components 1 and 2. The organisation of these genes and their vectors along the PCA mimics PS development; epiblast markers in the left quadrant (e.g. *Otx2* and *Fgf5*), early streak-related genes in the upper one (e.g. *Eomes*, *Mixl1* and *Foxa2*) and late streak markers further to the top-right (*Tbxt*, *Cdx2* and *Wnt3a*). This temporal sequence along dimension 1 ends with markers characteristic of caudal epiblast neural and mesodermal derivatives during early axial elongation (e.g. *Meox1*, *Pax3* and *Nr2f2*), reflecting the positioning of the distinct gastruloid samples in the PCA. The 48h and 72h DMSO replicates/conditions cluster in the lower left quadrant, followed by the 72h ActA and 72h low (0,5µM) CHIR samples. The 72h standard CHIR (3µM) replicates/conditions cluster further along dimension 1, suggesting they resemble a later PS developmental timepoint. Beyond this point, there is a bifurcation, with ActA gastruloids following a different trajectory in the PCA than that CHIR-treated gastruloids. **c**, Heatmap of gene expression (mean of the replicates variance-stabilised read counts) comparing 72h ActA and CHIR-treated gastruloids, with 48h and 72h DMSO gastruloids as a reference. ActA gastruloids exhibit a gene expression characteristic of the early PS, with higher levels of *Nodal*, *Cripto* (*Tdgf1*), *Eomes* and *Foxa2* and lower levels of *Tbxt*, *Cdx2*, *Wnt3a* in comparison to CHIR-treated gastruloids. Also, ActA gastruloids express higher levels of genes associated with the early epiblast (e.g. *Otx2* and *Fgf5*). The epiblast state of ActA gastruloids at 72h is more mature/advanced than that of 48h and 72h DMSO gastruloids because of the higher levels of *Nanog* and *Fgf5*, together with lower levels of *Utf1*. The 72h 0.5µM CHIR gastruloids display a gene profile more similar to ActA-treated gastruloids than 3µM CHIR gastruloids (e.g. *Eomes*, *Otx2*, *Myc* and *Cdx2*). The activation of the late PS module in CHIR gastruloids occurs in a concentration-dependent manner (e.g. *Cdx2*, *Wnt3a* and *Hoxa1*). **d,** Gene expression heatmap (mean of the replicates variance-stabilized read counts) of 96h CHIR (3µM) and ActA gastruloids. Gastruloids made with CHIR express genes that are associated with the early caudal epiblast (e.g. *Gdf11*, *Cdx2* and *Wnt3a*, with lack of *Wnt3*) and nascent mesoderm (e.g. *Tbx6*, *Cited1* and *Msgn1*). In contrast, ActA-treated gastruloids are delayed and express genes associated with the early to mid-streak transition (e.g. *Eomes*, *Mixl1*, *Wnt3* and *Cdx2*, and the lack of *Gdf11*). ActA gastruloids also show signs of having derivatives of the early PS as highlighted by the expression of *Noto* and *Gsc* (suggestive of node/notochord), *Foxa2* and *Sox17* (endoderm), *Kdr* and *Gata6* (haematoendothelium) and *Tbx6* and *Mesp1* (mesoderm). **e**, Euclidean distance analysis between gastruloid and embryo (pseudo-bulk atlas^86^, see **Methods**) datasets shows that both types of gastruloids follow a similar developmental progression. However, ActA gastruloids seem to resemble earlier developmental time points compared to CHIR gastruloids. This difference starts at 72h, with ActA gastruloids being more similar to E6.75 mouse embryos while CHIR-treated gastruloids resemble E7.25 embryos. **f**, HCR (MIP) on 72h ActA and CHIR gastruloids showing the expression of *Tbxt* and *Eomes* (in green and magenta, respectively). CHIR-treated gastruloids express only *Tbxt*, resembling the late posterior PS. In contrast, ActA gastruloids mimic the early PS, displaying both *Eomes* and *Tbxt* expression (white/greyish colour reflects colocalization). MIPs shown here are a representative selection of a total of 8 gastruloids per condition, from two independent experiments. The scale bar is equivalent to 100µm. **g,** HCR (MIP) highlighting the expression of PS and early differentiation marker genes in 96h gastruloids – total of 6 to 8 gastruloids imaged per condition, from two independent experiments. ActA gastruloids display a midstreak-like signature with high levels of *Tbxt* and expression of *Eomes* and *Wnt3*. On the contrary, gastruloids made with CHIR show weak expression of *Eomes* and *Wnt3* (brown and red arrowheads) and a reduced, polarized *Tbxt* expression. Similarly, genes related to endoderm, node/notochord and haematoendothelium are expressed in ActA gastruloids but show limited (yellow and orange arrowhead) or no expression in CHIR-treated gastruloids. Nascent mesoderm is present in both types of gastruloids as indicated by the expression of *Mesp1* and *Tbx6*. The scale bar is equivalent to 100µm.

These results suggest that, in gastruloids, Nodal signalling might trigger the early PS program. To further investigate the role of Nodal signalling in the PS and understand how it drives divergence in gastruloid development compared to the pulse of Wnt signalling, we treated gastruloids with ActA or CHIR and conducted a temporal comparative bulk transcriptomic analysis. PCA analysis indicates that ActA-treated gastruloids at 72h are more similar to gastruloids made with low CHIR (0.5μM) than to those treated with the standard CHIR dose (3μM) (**Fig. 3b**). This similarity was evident when examining key epiblast marker genes: ActA-treated gastruloids express high levels of both *Fgf5* and *Otx2*, low CHIR-treated gastruloids exhibit intermediate levels of these genes and 3μM CHIR-treated gastruloids express only *Fgf5* (**Fig. 3c**). Interestingly, these changes in the patterns of gene expression from high Nodal activity to increasing levels of Wnt signalling in gastruloids mirror the temporal progression of PS development during gastrulation. Gastruloids treated with ActA display an early PS-like signature (*Eomes*, *Foxa2* and *Tbxt* but no *Cdx2* or *Wnt3a*), whereas standard CHIR gastruloids mimic the late PS (no *Eomes* and *Foxa2*, but expression of *Tbxt*, *Cdx2* and *Wnt3a*) and low CHIR elicits an intermediate profile (low levels of *Eomes* and *Cdx2*, moderate *Tbxt* and absence of *Foxa2* and *Wnt3a)* (**Fig. 3c**). These results indicate that the Nodal pulse mediated by ActA triggers an early PS-like program in gastruloids that, although distinct from the standard CHIR one, shares similarities with that triggered by a low CHIR treatment.

The temporal differences between the two PS programs, as represented in ActA and CHIR gastruloids were further confirmed through Euclidean distance analysis: 72h ActA-treated gastruloids are more similar to E6.75 than 72h standard CHIR gastruloids, which closely resemble E7.25 mouse embryos (**Fig. 3e**). At later stages, PCA indicates a divergence in the development of the two types of gastruloids (**Fig. 3b**). At 96 and 120h, ActA-treated gastruloids express high levels of genes related to the specification and development of definitive endoderm e.g. *Foxa2*, *Cdh1*, *Epcam*, *Sox17* and *Cpm*^3,45^, node/notochord e.g. *Noto*, *Gsc*, *Foxa2* and *Shh*^46,47^, haematoendothelium e.g. *Kdr*, *Cd44*, *Flt1* and *Etv2*^5,48^ and anterior mesoderm e.g. *Tbx1*, *Gata6* and *Isl1*^6,49^ (**Fig. 3** and **Extended Data Fig. 7**). This anterior bias of ActA-treated gastruloids follows a highly anterior pattern of *Hox* gene expression (**Extended Data Fig. 7c**) and contrasts with the previously reported posterior character of gastruloids engineered with CHIR^24,50^, which develop essentially posterior neural and mesodermal-like structures (**Extended Data Fig. 7**). Concomitantly, at a phenotypical level, we observed that 120h ActA-treated gastruloids exhibit limited elongation characteristic of the gastruloids developed using CHIR (**Extended Data Fig. 7**), which would be expected in a representation of the anterior domain of the body plan. Noteworthy, these results were also noticed in gastruloids developed in distinct laboratories, using a different WT cell line, N2B27 media and culture conditions (**Extended Data Fig. 8** and **Methods**). Together, this indicates that distinct signalling activities play a key role in regulating the early and late PS and that a temporal progression from Nodal to Wnt signalling activity likely coordinates the emergence of the different anterior-posterior (AP) body structures.

### A temporal gradient of Nodal signalling activity controls the formation of anterior body structures in mammals

In our standard experimental conditions, while CHIR gastruloids stop developing around 120h, ActA-treated gastruloids develop at least until 144h. At this stage, these gastruloids continue to exhibit a round shape, with only around 50% showing some signs of elongation (**Fig. 4**). Interestingly, reducing the amount of ActA (from 100ng/ml to 25ng/ml) resulted in gastruloids displaying a significant elongation and a less pronounced anterior compartment (**Fig. 5**). Gene expression analysis revealed that genes highly expressed throughout the gastruloids treated with high ActA were localised to a small, anterior region in gastruloids treated with low ActA (e.g. *Kdr* and *Otx2*) (**Fig. 4** and **5**). Also, genes related to some anterior fates, such as *Nkx2-5* for cardiac mesoderm, were absent in low ActA gastruloids (**Fig. 5d**). In particular, we observed that in contrast to gastruloids treated with high levels of ActA, those treated with low levels developed a notochord-like structure with a tip that is positive for both *Tbxt* and *Noto*, indicating the presence of notochord progenitors (**Fig. 5d**). These results suggest that a low ActA treatment in gastruloids possibly results in the formation of more posterior body structures in comparison to high ActA.

**Fig 4.**
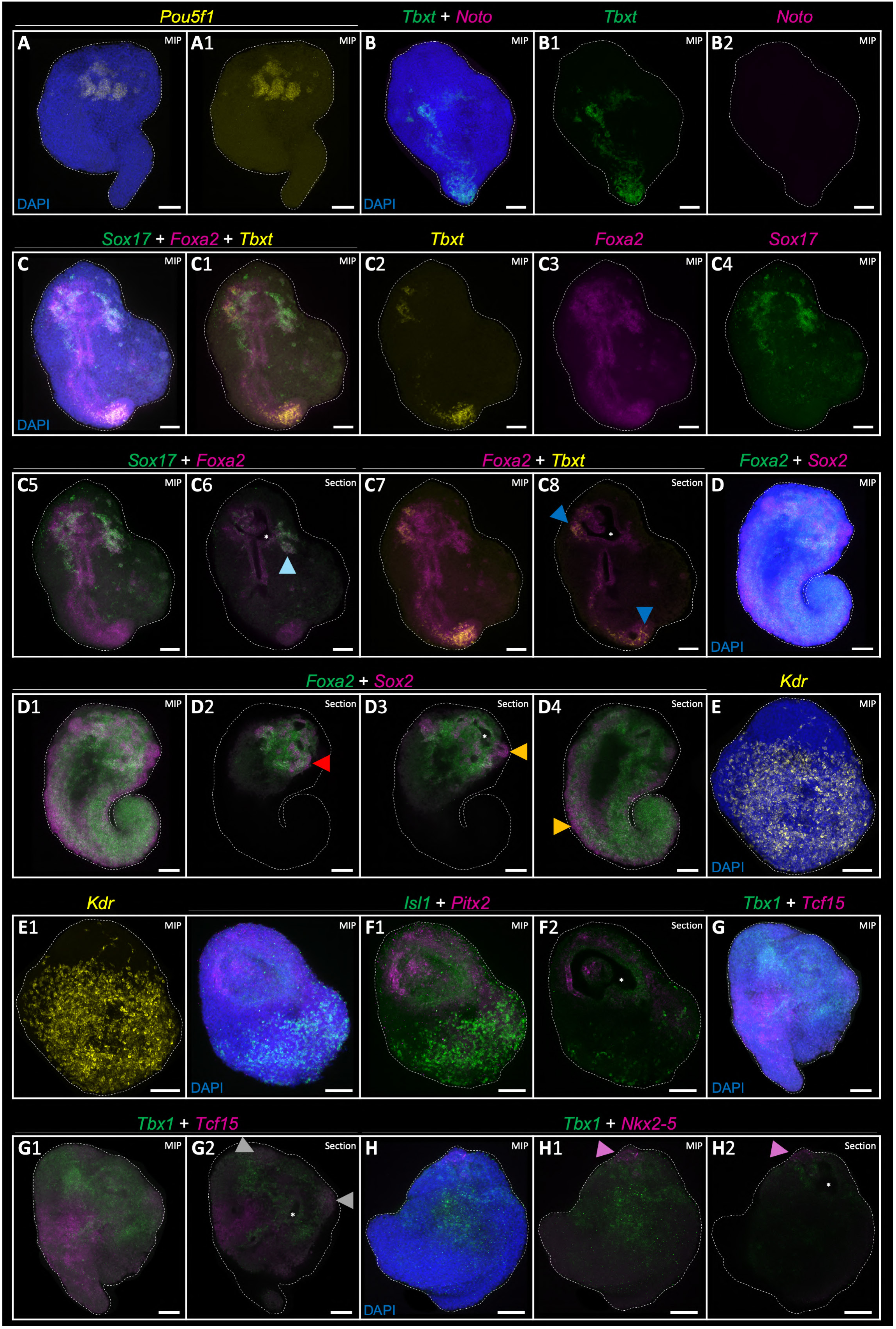
Activin A treated gastruloids develop several anterior body features. HCR images (MIP or optical section) for key genes involved in anterior body formation in ActA-treated gastruloids at 144h. 6-9 gastruloids were imaged per condition, four independent experiments. A – The localised *Pou5f1/Oct4* expression pattern is suggestive of PGC-like cells or ectopic pluripotency. B-C – *Tbxt* and *Foxa2* co-localisation indicates the presence of notochord-like cells, resembling the anterior-most part of the notochord in mouse embryos. This is inferred from the absence of *Noto*, a marker of notochord progenitor cells. The *Foxa2* expression that does not overlap with *Tbxt* suggests the presence of definitive endoderm-like cells. These cells show an organisation similar to the embryonic gut, including an internal cavity (white asterisk). Additionally, the co-localisation of *Foxa2* and *Sox17* at one end of this gut-like structure (in white/greyish) suggests its resemblance to the foregut of mouse embryos. D – Colocalization of *Foxa2* and *Sox2* (red arrowhead) is also suggestive of the presence of endoderm-like cells. However, *Sox2* expression alone (orange arrowhead) indicates the presence of neural tube-like cells, which can be observed going from one tip to the other, aligning with the gut and notochord as it occurs in the embryo. E-F – Endothelial-like cells are also present in ActA gastruloids, as shown by *Kdr* expression. *Isl1* and *Pitx2* expression patterns would be consistent with the development of pharyngeal endoderm and cardiac-like cells. G-H – *Tbx1* expression suggests the presence of anterior mesoderm. The co-localisation of *Tbx1* and *Tcf15* (grey arrowhead) suggests the presence of cranial mesoderm-like cells, while *Tcf15* expression adjacent to *Tbx1* is supportive of the existence of paraxial mesoderm-like cells. *Nkx2-5* expression near *Tbx1* suggests the presence of cardiac-like cells (pink arrowhead). The expression of this gene was only detected in around 75% of the gastruloids. The scale bar is equivalent to 100µm.

**Fig 5.**
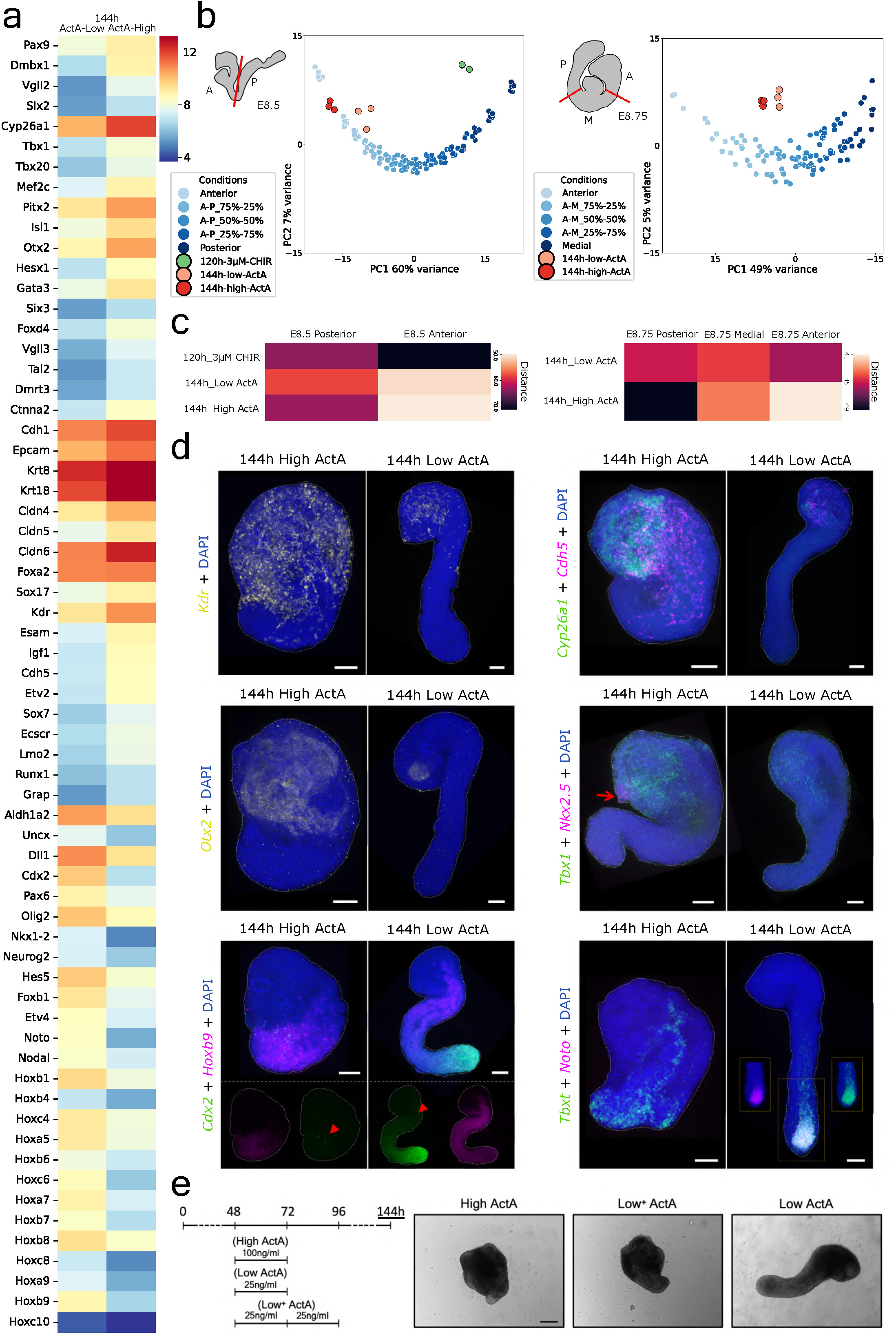
Nodal signalling activity drives anterior body formation in a dose/time-dependent manner. **a,** Heatmap showing gene expression (mean of the replicates variance-stabilised read counts; bulk RNAseq – three replicates per condition, from three independent experiments) in 144h gastruloids treated with varying levels of ActA. Gastruloids treated with higher doses of ActA exhibit increased expression of *Cyp26a1*, *Sox17*, *Pitx2*, *Pax9*, *Isl1*, and *Tbx1*, indicative of more anterior endoderm and mesoderm-like structures compared to gastruloids exposed to lower levels of Nodal signalling. In the latter gastruloids, *Aldh1a2/Raldh2*, which is typically found in the trunk region, is highly expressed along with a more posterior *Hox* gene signature (e.g. *Hoxa5* and *Hoxb8*). *Noto* expression, marking notochord progenitors, is limited to gastruloids treated with low ActA levels. Additionally, *Pax6* and *Olig2* are expressed at higher levels in low ActA gastruloids, suggesting the presence of more posterior neural tube-like cells compared to high ActA gastruloids. Increased expression of *Kdr*, *Esam* and *Cdh5* in gastruloids treated with a high dose of ActA suggests the presence of more endothelial-like cells in comparison to low ActA gastruloids. **b**, Comparative transcriptome analysis between bulk RNA-seq from different types of gastruloid and pseudo-bulk mixtures of E8.5 and E8.75 mouse embryos^56^ (see **Methods**), dissected in accordance to the schematics: E8.5 data was divided into “A” (anterior) and “P” (posterior) regions, whereas E8.75 data divided into “A” (anterior), “M” (medial) and “P” (posterior) compartments. PCA of bulk RNAseq data from gastruloid samples/replicates, projected onto computationally reconstructed E8.5 regional cell populations, suggest that 120h CHIR gastruloids cluster near the posterior region, while 144h gastruloids treated with high ActA cluster near the anterior region, though not fully resembling 100% anterior cell populations (“Anterior”). Low ActA gastruloids cluster closer to mixed anterior-posterior populations, suggesting they mimic more posterior regions of the body axis compared to gastruloids treated with high ActA. A similar analysis between high and low ActA gastruloids and E8.75 anterior and medial cell populations suggests that those treated with high ActA resemble more anterior embryo cell populations. **c**, Euclidean distance analysis between gastruloid and embryo RNAseq data^56^ (see **Methods** for more details). 120h CHIR-treated gastruloids differ from the anterior part of E8.5 embryos, while 144h ActA-treated gastruloids more closely resemble this region. Comparison with E8.75 embryo data suggests that 144h gastruloids treated with a higher ActA concentration mimic more anterior embryonic regions (anterior and medial) than those treated with lower doses, which display a broader body axis regional diversity. **d**, HCR representative MIPs of 4 independent experiments (6-8 gastruloids imaged per condition) highlighting differences between gastruloids treated with low and high ActA. *Kdr*, *Cyp26a1*, *Cdh5* and *Otx2* are widely expressed in high ActA gastruloids, whereas their expression is restricted to smaller regions in low ActA-treated gastruloids. *Tbx1* expression is higher and broader in high ActA gastruloids. *Nkx2-5* is also expressed in gastruloids developed with a high dose of ActA (red arrow) but absent in low ActA gastruloids. The expression of *Cdx2* is higher and broader in low ActA gastruloids (red arrowhead). *Hoxb9* expression is also broader in low ActA gastruloids. Co-expression of *Noto* and *Tbxt*, marking notochord progenitors, is observed only at the posterior tip of gastruloids treated with a low dose of ActA. Inlets highlight individual gene expression. The scale bar is equivalent to 100µm. **e**, Representative images of gastruloids (three independent experiments) highlighting the temporal effect of prolonged exposure to Nodal signalling activity (see also **Extended Data Fig. 10**). 144h gastruloids treated with low levels of ActA between 48 and 96h are morphologically similar to standard high ActA gastruloids (treatment between 48h and 72h) and display minimal signs of elongation, which contrasts with standard low ActA gastruloids. The scale bar is equivalent to 50µm.

To investigate the importance of the levels of Nodal signalling in PS and further clarify the anterior-posterior (AP) differences in body axis formation between gastruloids treated with high and low levels of ActA, we performed bulk RNAseq at multiple time points. Unlike gastruloids exposed to different levels of CHIR, gastruloids treated with high or low concentrations of ActA did not exhibit many significant differences at 72h (**Extended Data Fig. 9**). While we observed slight changes in the levels of *Tbxt*, *Nodal*, *Eomes*, *Pitx2*, *Fgf8* and *Tdgf1*/*Cripto*, there were no significant changes in the expression of Nodal targets like *Lefty1/2* (**Extended Data Fig. 9**). The main difference we observed concerns *Foxa2*, a critical mesendodermal transcription factor ^51,52^, whose expression was upregulated in gastruloids with high levels of ActA (**Extended Data Fig. 9**). At 96h, we noticed differences in both the activity of Nodal signalling (higher levels of ActA resulted in more expression of *Cer1* and *Lefty1/2*) and the expression of key PS-related genes between both types of ActA-treated gastruloids (**Extended Data Fig. 9**). On the one hand, gastruloids treated with high levels of ActA exhibit stronger expression of genes associated with the early PS, like *Lhx1*, *Cripto*, *Foxa2* and *Gsc*. On the other hand, gastruloids treated with lower levels display a significant expression of genes associated with the late PS to CLE transition, such as *Wnt3a*, *Wnt8a*, *Cdx1/2* and *Nkx1-2* (**Extended Data Fig. 9**). Importantly, these differences seem to impact fate specification as we noticed differences in the expression of genes associated with, for instance, extra-embryonic mesoderm^53^ and endoderm (e.g. *Flt1* and *Sox17*, respectively) (**Extended Data Fig. 9**).

A thorough transcriptomic evaluation between the two types of ActA gastruloids at 144h indicates differences not only in the amount of anterior tissues but also in their nature. For instance, higher levels of *Pax9* and *Sox17*, together with *Foxa2*, suggest the presence of pharyngeal endoderm ^54,55^ in gastruloids treated with high levels of ActA but not in those treated with low levels of this Nodal surrogate. The endothelial gene expression signature in high ActA gastruloids is more complete, being positive for *Kdr*, *Esam*, *Igf1*, *Cdh5* and *Etv2*, in contrast to only significant levels of *Kdr* being detected in low ActA-treated gastruloids (**Fig. 5a**). Also, higher levels of *Tbx1*, *Pitx2* and *Isl1* are consistent with the development of more anterior mesoderm-like tissues in gastruloids treated with a higher dose of ActA. The higher expression of *Hesx1*, *Dmbx1* and *Cyp26a1* in these gastruloids, in contrast to higher levels of *Raldh2*, *Pax6* and *Foxb1* in low ActA gastruloids (**Fig. 5a**), further suggests the development of distinct AP body structures in a concentration-dependent manner, with low ActA-treated gastruloids developing more posterior ones, spanning broader AP body axis regions (**Fig. 5**). This can be noticed by a more posterior Hox gene expression (e.g. positive for *Hoxb9*) in gastruloids treated with low levels of ActA (**Fig. 5a** and **Extended Data Fig. 7**) and is supported by two complementary bioinformatic analyses comparing the two types of ActA gastruloids and embryonic data^56^ obtained from different regions of E8.5 and E8.75 mouse embryos (**Fig. 5b** and **Methods**).

These results suggest two key points: 1) a temporal decreasing gradient of Nodal signalling in the PS may be essential for establishing the full AP diversity of anterior mesendodermal tissues and 2) there is likely an inverse relationship between Nodal and Wnt signalling in gastruloids. To better understand the former, we developed low and high ActA gastruloids with varying durations of the treatment and noticed that there is a relationship between the two variables, with low ActA gastruloids treated between 48h to 96h exhibiting a morphology more akin to that of high ActA gastruloids that were only treated between 48h and 72h (**Fig. 5e** and **Extended Data Fig. 1**0). These observations provide a rationale for how the varying temporal levels of Nodal signalling activity in PS (from high to low) influence the formation of different AP mesendodermal body structures. To understand how this relates to the dynamics of Wnt signalling, we evaluated both Wnt and Nodal reporter activity in gastruloids treated with different doses of ActA (**Fig. 6**). At 72h, high levels of ActA elicit a robust Nodal signalling response with low Wnt signalling that, in most cases, disappears with time. In contrast, low levels of ActA resulted in higher levels of Wnt signalling at 72h. By 120h, the majority of these gastruloids treated with low ActA exhibited high levels of Wnt signalling at the posterior part and some Nodal activity in the anterior compartment.

**Fig 6.**
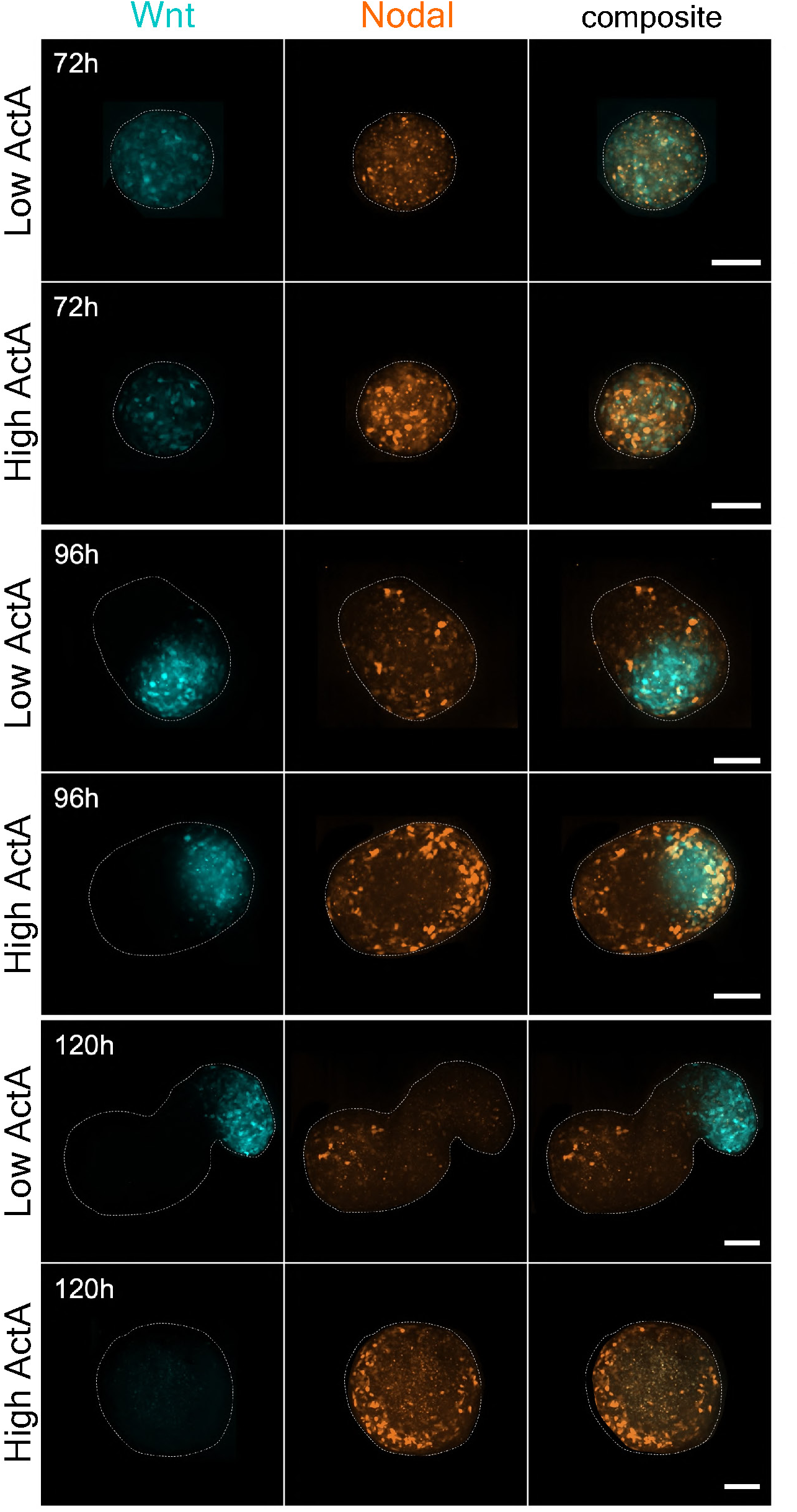
Nodal and Wnt signalling activity in gastruloids treated with different levels of ActA. At 72h, higher levels of Nodal activity (orange) are observed when a higher dose of ActA was used to make gastruloids. In contrast, Wnt signalling (cyan) is more active in low ActA gastruloids. Both types of ActA-treated gastruloids display a polarised domain of Wnt activity at 96h. Nodal activity is increased in gastruloids treated with high levels of ActA and is more prominent in the outer part of the gastruloids. At 120h, we observed a mixture of elongated and round gastruloids. Elongated gastruloids were more frequently observed in the low ActA condition and they showed strong Wnt levels posteriorly and some Nodal activity in the anterior compartment. Non-elongated gastruloids were mostly observed in the high ActA condition and they exhibited significant Nodal activity in the outer shell, more pronounced towards one side of the gastruloid, and low levels of Wnt signalling. Scale bar is equivalent to 100µm.

Together, these observations provide evidence for how different levels of Nodal signalling control Wnt activity and suggest the existence of an antagonistic interaction between Nodal and Wnt signalling in the PS (**Fig. 7**). Overall, this study provides mechanistic insights into how the spatiotemporal activities of Nodal and Wnt signalling regulate fate decisions in the PS by triggering two distinct developmental modules, governing the emergence of the early mammalian body plan. Also, in varying the nature, levels and duration of the signalling treatments provided at 48h, we showed that gastruloids can, independently, mimic the two main PS modules.

**Fig 7.**
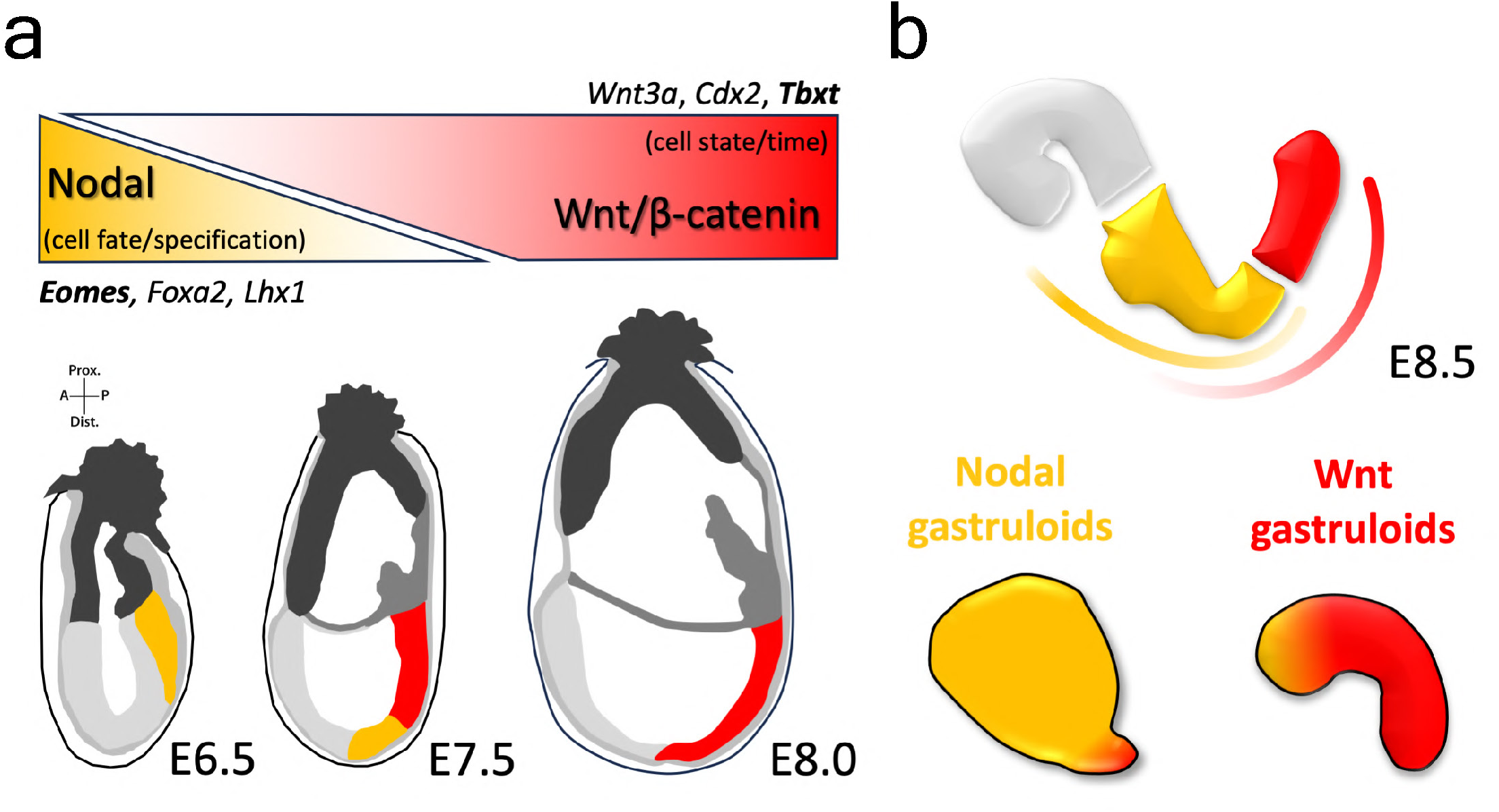
A temporal coordination between Nodal and Wnt signalling governs the emergence of the mammalian body plan. **a,** Schematic representation of the signalling-dependent modularity in the mouse PS during gastrulation. The early PS (in yellow) is mostly controlled by Nodal signalling, whose activity decreases over time and is responsible for setting out the various anterior mesendodermal fates through the activation of key genes like *Eomes*, *Foxa2* and *Lhx1*. Conversely, Wnt/β-catenin activity increases over time and regulates the late posterior PS (in red) and the emergence of the CLE, through the activation of critical genes such as *Tbxt*, *Cdx2* and *Wnt3a*, thus pacing gastrulation. Extra-embryonic tissues and other non-PS embryonic tissues are shown using distinct shades of grey. **b**, Schematic representation of an E8.5 mouse embryo highlighting the two compartments that arise from the early and late PS modules and how they are regulated by the opposing gradients of Nodal and Wnt signalling activity in the PS (in yellow and red, respectively). Brain tissues (in blue) relate to a distinct gastrulation module, which requires the absence of Wnt and Nodal signalling activity. A comparative representation of gastruloids treated with Nodal/ActA or Wnt/CHIR is also made using the respective colours, in a gradient manner.

## Discussion

Current views of gastrulation in mammals suggest that it is a continuous process driven by a cooperation between Nodal and Wnt signalling in which cells are allocated fates orderly from anterior to posterior^4,9,57^. Recent studies have shown that two T-box transcription factor modules divide the PS into two defined spatiotemporal compartments. One, dependent on *Eomes* establishes fates in the early/anterior PS and a second, later acting, governed by *Tbxt*, promotes posterior fates^16,58–60^. Here using gastruloids (and see also Wehmeyer et al.^61^) as a model system for mammalian gastrulation, we show that each of these transcriptional modules is controlled by a specific signal: Nodal for *Eomes* and Wnt for *Tbxt*, and that the order and pace of gastrulation relies on opposing cross-regulatory interactions between the signalling pathways (**Fig. 7**). In this process, our results suggest that Wnt and Nodal have different roles.

The role of Wnt signalling during gastrulation has sparked various ideas about the relationship between time and space in cell fate assignments. The two prevailing hypotheses suggest that Wnt signalling drives mesoderm specification or acts as a “posteriorizing agent”^40,62^. Our findings support the latter hypothesis as changes in the levels of Wnt signalling did not alter the relative amounts of specific fates, in particular mesoderm. However, as a posteriorizing agent rather than acting as a morphogen in a classical sense^62^, our results suggest that Wnt/β-catenin signalling works mainly as a timer, controlling the pace of gastrulation and the ordered emergence of axial progenitors. Furthermore, our results suggest that this process is gradual and involves, in the first instance, the maturation of early epiblast cells towards a Nanog-positive posterior epiblast pluripotent state, and the concomitant inhibition of neural differentiation. This is in line with genetic studies performed in the mouse embryo evaluating the importance of the Wnt-related T-box transcription factors *Eomes* and *Tbxt* in suppressing pluripotency and neural differentiation^42^, and with *β-catenin* mutant embryos lacking the characteristic posterior *Nanog* expression at E6.5^41^. The emergence of multipotency and axial elongation occurs through the activation of the late PS, contingent on *Tbxt*, in a Wnt dose/time-dependent manner. This “posteriorization” effect is associated with an acceleration of the Hox timer, whose activation and control have been shown to depend on Wnt signalling^50,63,64^, and a critical reduction of Nodal signalling, which decays during gastrulation as the levels of Wnt increase and enable the formation of posterior body structures^20,21^.

In contrast to the response of cells in the gastruloid to varying concentrations of CHIR, changing the levels of Nodal signalling affects cell fates within the anterior mesendodermal compartment. This result is consistent with studies on the dose-dependent effects of Nodal/Smad signalling during mouse gastrulation^65^ and work on amphibian animal cap explants treated with ActA^66^. In gastruloids, we observed that while high and prolonged levels of ActA elicit the most anterior fates (pharyngeal endoderm and cardiac/cranial mesoderm), low/brief signalling activity results in a broader anterior-posterior distribution of fates and even the expression of some posterior *Hox* genes. Here, we noticed that Nodal signalling inversely affects Wnt activity (see also Wehmeyer et al.^61^ and McNamara et al.^67^) and therefore, this *Hox* gene expression signature in gastruloids treated with low levels of ActA is probably due to the release of some Wnt signalling that is suppressed by high levels of Nodal signalling. The molecular details of the Nodal effects on Wnt signalling remain an open question but it is likely that, together with the reciprocal effects of Wnt on Nodal signalling, they govern the emergence of the mammalian body plan.

Gastruloids are a versatile tool to explore various features of mammalian development ^68^. However, a critical element of these studies is the interpretation of the model, in particular its relationship to the embryo. It is now clear that while CHIR driven standard mouse gastruloids display some anterior mesendodermal elements emerging sporadically in different experiments, they mostly represent the posterior part of the mouse embryo, with derivatives from the CLE^22,24,26–28,69–71^. The work presented here and in Wehmeyer et al.^61^ supports this assignment and provides a rationale for how it occurs.

Our results show that at 48h after aggregation, most cells have exited the naïve pluripotency state and begin implementing the gastrulation programme with the expression of *Nodal*, *Wnt3 and Eomes*. The degree to which this happens in individual gastruloids depends on the cell line and culture conditions. In raising the levels of Wnt signalling, the addition of CHIR triggers a genetic programme resembling the late PS, involving *Tbxt*, *Wnt3a*, *Cdx2* and *Hox* genes, leads to the suppression of the anterior module and ultimately results in the development of a CLE-like neuromesodermal competent cell population that generates spinal cord and paraxial mesoderm derivatives. The amount and degree of anterior fates in CHIR-treated gastruloids will most likely depend on the levels of Nodal signalling accrued by the cells at around 48h as defined by cell line and the culture conditions.

Replacing CHIR by Activin A increases the levels of Nodal signalling and enhances the gastrulation programme, through the upregulation of key genes expressed in the early PS, such as *Eomes*, *Foxa2* and *Lhx1*. This results in the formation of mesendodermal-like structures that are related to different AP levels of the body, depending on the concentration and duration of the ActA treatment. Given these results and the known signalling dynamics of Wnt and Nodal in the mouse embryo, we propose that the antagonistic interaction of these signalling pathways in setting two distinct developmental modules in the PS plays a critical role in establishing the early mammalian body plan. Here, in agreement with previous suggestions^72^, Wnt signalling appears to coordinate these events by acting as a pacemaker for the process.

Together with the results of Wehmeyer et al.^61^, our results suggest that in the embryo, the interactions between Wnt and Nodal signalling over time modulate the outcome of the interactions between Eomes and Tbxt. This modulation results in the timely association of cell fates.

Finally, an obvious feature of gastruloids when compared with embryos is their lack of a structure similar to the PS. Our results (and see also Fiuza et al.^73^) suggest that the reason for this might be that gastruloids represent the PS and, therefore, reveal an hereto unknown self-organising ability of this structure. We foresee that this proposal for the nature of gastruloids and the work presented here sets the stage for researchers to use the signalling-dependent modularity of the PS and the ability to make gastruloid chimaeras in favour of the engineering of specific body structures in vitro with biomedical potential, given the ability to grow human gastruloids^74^.

## Acknowledgements

We thank Anna Bigas for the E14Tg2A cell line, Joshua Frenster for providing mEpiLSCs, Thomas Gregor for the Cdx2 antibody, Tina Balayo for management and cell culture assistance, Raquel Flores Peirats for management assistance, Veronika Mantziou and Tina Balayo for the optimisation of the HCR protocol, Soyeon Park for assistance with RT-qPCRs, the laboratories of Jared Toettcher and Alfonso Martinez Arias, particularly Ulla-Maj Fiuza and Sara Bonavia, for useful discussions. We would also like to thank the IGC/GIMM Genomics Unit, the CRG Microscope facility and the UPF/CRG Flow Cytometry Unit. This work was supported by an EMBO Postdoctoral Fellowship (ALTF 948-2022) to AD; a Generalitat de Catalunya AGAUR Grant (2021 SGR 00175) to AD and GTC; a PhD fellowship (FPU18/05091) from the Spanish Ministry of Science, Innovation and Universities to GTC; a PhD fellowship from Boehringer Ingelheim Fonds to SB; the project PID2021-127311NB-I00 financed by the Spanish Ministry of Science and Innovation, the Spanish State Research Agency, the European Regional Development Fund (FEDER) and the ICREA Academia program to PCG and JGO; the Lewis-Sigler Scholars program to HMM; the Swiss National Science Foundation grant (SNSF 407940_206405) and the Human Frontier Science Program (HFSP LT000032/2019-L) to AM; the German Research Foundation (DFG) project grants AR 732/3-1, AR 732/2-1, CRC 992 (project ID 192904750) and Germany’s Excellence Strategy grant (CIBSS – EXC-2189 – project ID 390939984) and the Ministry of Science and Education of the State of Baden Württemberg to SJA; and an ERC Advanced Grant (834580) to AMA. Research at the AMA lab was also funded by the Maria de Maeztu Program for Units of Excellence in R&D (CEX2018-000792-M).

## Contributions

AD and AMA conceived the project and interpret the results; AD designed the experiments and performed the E14Tg2A and TLC-mCherry E14 gastruloid work, HCRs and bulk RNAseq; PPM and GTC processed and, together with AD, analysed the transcriptomic data; GR and AD made and characterised the Wnt3 mutant gastruloids; SB and AD did the work shown as IF stainings; PCG performed 3D segmentation and quantification of IF stainings and, together with AD, SB, JGO and AMA, analysed the data; HMM did the Wnt/Nodal double reporter work; YR and AM provided 129/svev gastruloids; SS and MV performed preliminary experiments using P300 proximity labelling in CHIR and ActA gastruloids. SJA and AW shared unpublished data and provided essential input for the interpretation of the results; AD wrote the original draft; AMA reviewed and edited the manuscript; All authors commented on the manuscript.

## Methods

### Cell lines and culture

In this work we used wildtype (WT) *E14Tg2A* mESCs obtained from the lab of Anna Bigas at the Hospital del Mar Research Institute (Spain), WT *129/svev* from EmbryoMax© (CMTI-1), the Wnt TCF/Lef reporter cell line “*TLC-mCherry*^21,75^ and a Wnt and Nodal double reporter cell line^67^. EpiLSCs originate from the *E14Tg2A* mESCs and were developed in accordance with Joshi et al.^76^. The *Wnt3*KO cell line was made using E14Tg2A mESCs (see below).

The WT and *Wnt3*^-/-^ *E14Tg2A* and *TLC-mCherry* cell lines were cultured in ESLIF media (10% fetal bovine serum; ThermoFisher, 10270106), on 0.1% gelatine in 1X Phosphate Buffered Saline (PBS; Biowest, X0520) coated 6-well plates (Greiner, 657160) as described in Baillie-Johnson et al.^77^. L-Glutamine (ThermoFisher, 25030024) was used instead of Glutamax. These cells were cultured in a humidified incubator at 37°C with 5% CO2 and passaged every three days, with a seeding density of approximately 6,5x10^4^ cells per well. The double reporter cell line was cultured in agreement with McNamara et al.^67^, but ESLIF was used instead of 2iLIF. 129/svev mESCs were cultured as described in Mayran et al.^26^, using a DMEM based media supplemented with 10%FBS and 2iLIF. All cell lines were maintained in culture for at least two to three passages post-thawing before experimental use and were routinely tested for mycoplasma.

### *Wnt3* mutation

Oligos used as single guide RNA (sgRNA; see **Supplementary Table 1**), targeting the 5’ and the 3’ region of exon 2 of the *Wnt3* gene, were annealed (94C for 4 min, 70C for 10 min and 37C for 20 min) and cloned into the pSpCas9(BB)-2A-GFP plasmid (Addgene, #48138)^78^. The cloning was done through a Golden Gate reaction and using the BbsI enzyme (New England Biolabs, R3539S). *E14Tg2A* mESCs were plated on 6-well plates at a density of 150K/well one day before lipofection and transfected with the assembled Cas9-gRNA plasmids at equimolar ratios using Lipofectamine 3000 (Thermo Fisher Scientific, L3000001), according to the manufacturer instructions. One week post-transfection, GFP-expressing cells were sorted via fluorescence-activated cell sorting (FACS; BD Influx™ Cell Sorter) and single-cell distributed into gelatin-coated 96-well plates to obtain single, independent clones. From these clones, genomic DNA (gDNA) extraction was performed using the NZYTech tissue gDNA isolation kit (MB13502) and used for PCR validation and Sanger sequencing, using the primers described in **Supplementary Table 1**. Clone 29, with a deletion of exon 2, was chosen based on the morphological characterisation of Wnt3^-/-^ CHIR and DMSO treated gastruloids at 120h (see **Extended Data Fig. 5a**). Further genetic confirmation was done via reverse transcription-quantitative polymerase chain reaction (RT-qPCR; see below) and the results (**Extended Data Fig. 5b**) are in agreement with the literature^17,79^.

### Gastruloid engineering

*WT and Wnt3^-/-^ E14Tg2A* and *TLC-mCherry s*tandard (CHIR) gastruloids were generated in accordance with previously published work ^22,23,77^ using 300 cells as the starting number of cells, non-adherent U-bottom 96-well plates (Greiner, 650185) and NDiff®227 (N2B27) media from Takara (Y40002; batch references: ‘AM30020S’, ‘AM10016S’, ‘AM90020S’, ‘AN30020S’ and ‘AO30013S’). All N2B27 media was batch-tested before the experiments described in this study to assess their ability to generate at least 80% of CHIR-treated gastruloids and the degree of morphological inter- and intra-batch variation. In standard gastruloids, Dimethyl sulfoxide (DMSO; Sigma, D2438) was used as a vehicle for the CHIR treatment. DMSO gastruloids only contained DMSO and were used as a control for the amount of DMSO used in gastruloids treated with varying levels of CHIR. ActA gastruloids were made similarly to CHIR-treated gastruloids, but Activin A (High - 100ng/ml, Low - 25ng/ml; Recombinant human/mouse/rat Activin A PLUS protein, (Qkine, Qk005) was used instead of CHIR (Chi99021, Sigma, SML1046) between 48 and 72h. High^-^ ActA-treated gastruloids were developed using 100ng/ml of ActA between 48 and 72h but were washed two consecutive times with 150µl of fresh N2B27 media (without ActA) at the end of the treatment, before continuing the standard protocol. Low^+^ ActA gastruloids were developed similarly to low ActA gastruloids, but 25ng/ml of ActA was also added between 72h and 96h. AF-treated gastruloids were cultured with ActA (25 ng/ml) and bFGF (12.5 ng/ml; Recombinant Murine FGF-basic, Peprotech, 450-33) between 0 and 48h. ActA (Qkine) was stored following the instructions from the manufacturer (Qkine) and used fresh or 4°C maintained for a maximum of 48h. Nodal and Wnt double reporter gastruloids were made using 200 cells per gastruloid, following McNamara et al.^67^. 129/svev gastruloids were developed by plating 300 cells per gastruloid and cultured in homemade N2B27 media as described in Mayran et al^26^.

### Standard imaging and Wnt and Nodal reporter analysis

WT and *Wnt3*^-/-^ E14Tg2A brightfield gastruloid images were acquired using an Olympus brightfield CKX53 microscope (4x and 10x lens) equipped with an EP50 camera, and a Zeiss Cell Observer microscope (5x/0.16 or 10x/0.3 Ph lenses depending on the size of the gastruloid). *TLC-mCherry* gastruloids were imaged in a Zeiss Cell Observer fluorescence microscope, filter sets: DAPI/CFP/GFP/YFP/RFP/Cy5, using 10x/0.3 Ph or 20x/0.5 Ph lenses (depending on the size of the gastruloid). These images were processed using Fiji ^80^. Wnt and Nodal double reporter gastruloids were imaged and processed as in McNamara et al.^67^.

### Bulk RNA sequencing

RNA for bulk RNA sequencing was extracted from gastruloids using the Qiagen RNeasy Mini Kit (74104). The number of pooled gastruloids varied according to their developmental stage: full 96-well plate if at 48, 60h and 72h, half a 96-well plate if at 96 and one-third of a 96-well plate if at 120 or 144h (at least 3 replicate samples were obtained per condition, from several independent experiments). RNA was also extracted from 2D cultures of ESCs and EpiLSCs (T=0h) following the same procedure. RNA concentration and purity were determined using a Fragment Analyzer (Agilent). cDNA libraries (non-stranded) were prepared from total RNA using the SMART-Seq2 protocol^81^. Sequencing was performed on an Illumina NextSeq 2000 at the IGC Genomics Facility (Instituto Gulbenkian de Ciência, Portugal), generating around 25 million single-end 100 base pair reads per sample.

### Bulk RNA sequencing data preprocessing

The quality of the raw sequencing files was initially assessed using *FastQC* (version 0.11.9)^82^. Adapter trimming and low-quality filtering were performed using *TrimGalore!* (version 0.6.7)^83^ where reads of Phred score below 20 were removed. Following trimming, the quality of the cleaned reads was re-evaluated with *FastQC* as a quality checkpoint to ensure the effectiveness of the trimming and filtering steps.

The processed sequencing files were mapped to the mouse genome (GRCm38.p6) using the GENCODE annotation (version 25) with the *STAR* alignment algorithm (version 2.7.10a)^84^. The mapping was performed with the default settings and the aligned read counts were generated using the *quantMode* function within *STAR*. The resulting count matrix was created by concatenating the assembled reads using the *Python*3 environment.

### Bulk RNA sequencing analysis and visualisation

The downstream statistical and visualisation analysis was performed in a *Python* 3.10.12 environment. Gene expression counts were normalised using the *pyDESeq2* package (version 0.4.0). Variance-stabilising transformation (VST) was used to transform raw counts across samples^85^. Feature selection and dimensionality reduction using PCA were performed with *sklearn* (version 1.2.2), while *seaborn* (version 0.13.2) and *matplotlib* (version 3.7.2) were used for plotting. Gene differential expression analysis was also conducted using *pyDESeq2*.

Genes with an adjusted p-value < 0.1, as determined by the Benjamini-Hochberg method for multiple testing correction, and an absolute log2 fold change > 0.5 were determined as significantly differentially expressed (DEGs).

Mixed populations of embryo sections from Imaz-Rosshandler et al.^56^ and Pijuan-Sala et al.,^86^ were computed by creating random pseudo-samples using *KFold* from *sklearn*. A 5% step was set to create a gradient of mixed conditions.

### RT-qPCR

RNA for RT-pCRs was obtained as described above for bulk RNAseq, from WT and *Wnt3* mutant gastruloids treated with either 3µM CHIR or the corresponding amount of DMSO. Complementary (cDNA) synthesis from total RNA was performed using the SuperScript IV First Strand Synthesis System (Invitrogen) with random hexamer primers. Gene expression was quantified using LightCycler® 480 SYBR Green I Master (Roche) according to the manufacturer’s instructions, with 200nM primers and 12ng/μl cDNA in a 10μl final reaction volume. Thermocycling was conducted using a LightCycler® 480 (Roche) qPCR machine. The primers are listed in **Supplementary Table 1**. *Hmbs* and *Atp5pb* were used as house-keeping genes.

### In situ hybridisation chain reaction

The HCR protocol for gastruloids was adapted from Choi et al.^87^ and the HCR probes and amplifier hairpins were purchased from Molecular Instruments. Gastruloids were first fixed in 4% paraformaldehyde (PFA; Electron Microscopy Sciences, 15710) in PBS without calcium and magnesium (PBS^-/-^; Sigma, D8537), supplemented with 0.1% Tween-20 (Sigma, P9416; PBST) for 20 to 25 minutes at room temperature (RT), depending on the developmental stage of the gastruloids. Following two washes in PBST, the gastruloids were dehydrated in methanol (MeOH; series of 25% increase in PBST) and stored at -20°C. Prior to initiating the HCR procedure, gastruloids were rehydrated through a reverse MeOH/PBST series. After two washes in PBST, gastruloids were incubated in hybridisation buffer at 37°C for 30 minutes, and then overnight (ON) with the different gene probes. On the following day, gastruloids were washed four times every 15 minutes in 30% probe wash buffer and three times for 5 minutes each in 5X SSC buffer (Sigma, S6639) with 0.1% Tween-20 (SSCT). For the pre-amplification step, gastruloids were incubated in amplification buffer for 30 minutes at RT. The hairpins were snap-cooled (90 seconds at 95°C, followed by 30 minutes at RT) before being added to the amplification buffer solution where the gastruloids were incubated ON (RT). The next day, gastruloids were washed four times for 5 minutes each, plus three additional times every 20 minutes in SSCT (RT) before being stained with 4’,6-Diamidino-2-Phenylindole, Dihydrochloride (DAPI; ThermoFisher, D1306) ON at 4°C. Finally, the gastruloids were washed in PBST and mounted in depression slides (Sigma, BR475505) with RapiClear 1.47 (SunJinLab, RC147001) as described in Dias et al.^88^. Imaging was done on a Zeiss LSM980 Airyscan 2 microscope, with 405 nm, 488 nm, 561 nm and 639 nm laser filter sets, and using 10x/0.45 or 20x/0.8 lenses (depending on the size of the gastruloid). Z-stack (5-10µm) images were analysed using Fiji/ImageJ.

### Immunohistochemistry

*E14Tg2A* WT CHIR/DMSO and Wnt3 DMSO mutant gastruloids (standard treatment, four independent experiments) were prepared for IF stainings through fixation in 4% PFA in PBS^-/-^ during 2h at 4°C. After two washes in PBST, gastruloids were dehydrated through MeOH series in PBST (25% increase, every 5 minutes), until reaching 100% MeOH, and stored at -20°C until use. Before starting the IF staining per se, gastruloids were rehydrated through series of 25% decreasing MeOH concentrations at RT and washed two times in PBST. Samples were then permeabilised 3x10 min in PBST (with 0,5% Triton X-100) and blocked 1,5h in blocking buffer made of PBST with 10% bovine serum albumin (BSA, Merck Life Science, Cat#3117332001) at RT. Following this, gastruloids were incubated with primary antibodies (**Supplementary Table 2**) diluted in PBST with 2% BSA (i.e. wash buffer) for approximately 48h at 4°C, shaking. Samples were then rinsed 6x10 minutes in wash buffer at RT and incubated with secondary antibodies (**Supplementary Table 2**) with DAPI (1:5000), both diluted in wash buffer, for approximately 24h at 4°C with shaking. Following 6x10 minutes rinses in wash buffer at room temperature, samples were rinsed once in PBST and mounted in depression slides with RapiClear 1.47 as described in Dias et al.^88^. Imaging was done during the last day of the protocol on a Zeiss LSM980 Airyscan 2 microscope with 405 nm, 488 nm, 561 nm and 639 nm laser filter sets and a 20x/0.8 lens. A total of 142 gastruloids were imaged (from 18 independent IF experiments), with Z-stacks of 1024 × 1024 images acquired every 2μm. The laser intensities were kept stable throughout the entire experiment. All gastruloids were imaged with 1x digital zoom, except for those at 48h which enabled the use of 1.2x digital zoom. A total of 145 gastruloids were imaged, with an average of 4-5 gastruloids per condition.

### 3D image segmentation

3D cell segmentation was performed with QLIVECELL (https://github.com/dsb-lab/qlivecell; *Casaní-Galdón et al., in preparation*) using the StarDist 2D pretrained model *2d_versatile_fluo*^89–91^. Segmentation masks were classified as cells if they were present in at least two z-planes and if their size was above a threshold in the centre plane, defined as the z-plane with the highest summed fluorescence intensity. To minimise potential crosstalk between channels, the 2D centre of each mask was calculated as the centroid in this plane and the corresponding 3D coordinate was used as the cell centre. Size thresholds were computed per time point from the distribution of mask areas in the centre plane. Kernel density estimation (KDE) was applied to the histogram of mask areas, using the smallest kernel width yielding a single local minimum (**Supplementary Data Fig. 1**) - the minimum was used to separate true cells from segmented debris. Overall, segmentation successfully identified at least 75% of the cells in each gastruloid. The number of segmented cells can be found in **Supplementary Data Table 3**.

### 3D image analysis

To mitigate the influence of extreme values on downstream analysis, outliers were removed from the dataset using the interquartile range (IQR) method. The first (Q1) and third (Q3) quartiles were computed for each condition and marker, and the IQR was defined as the difference between Q3 and Q1. Data points falling below Q1 − 3.5×IQR or above Q3 + 3.5×IQR were classified as outliers and excluded from further analysis. This procedure was applied independently per condition and time.

The mean fluorescence per channel was computed using the segmented mask at the centre plane of each cell. To correct for inter-gastruloid technical variability, background subtraction was performed using the mean intensity in a 50×50 px region at the top-left corner of each image, per channel and z-plane.

Signal drift across the z-axis introduced artificial correlations between independent channels (**Supplementary Data Fig. 2b**). To correct for this effect, each z-plane was normalised according to its intensity profile and scaled by the overall mean (**Supplementary Data Fig. 2a**). Similar to the co-expression scatter plots, histograms showing both corrected and non-corrected signal intensities (**Fig. 2** and **Supplementary Data Fig. 3**, respectively) were made using data from pooled 3D gastruloids.

## Data Availability

Raw and processed RNA sequencing data were submitted to Biostudies/ArrayExpress and have the following accession information: “E-MTAB-15149”. The bioinformatics analysis pipeline, including all code and parameters, is available on the following GitHubs: https://github.com/stembryo-lab and https://github.com/dsb-lab/qlivecell).

---

**Extended Data Fig 1.**
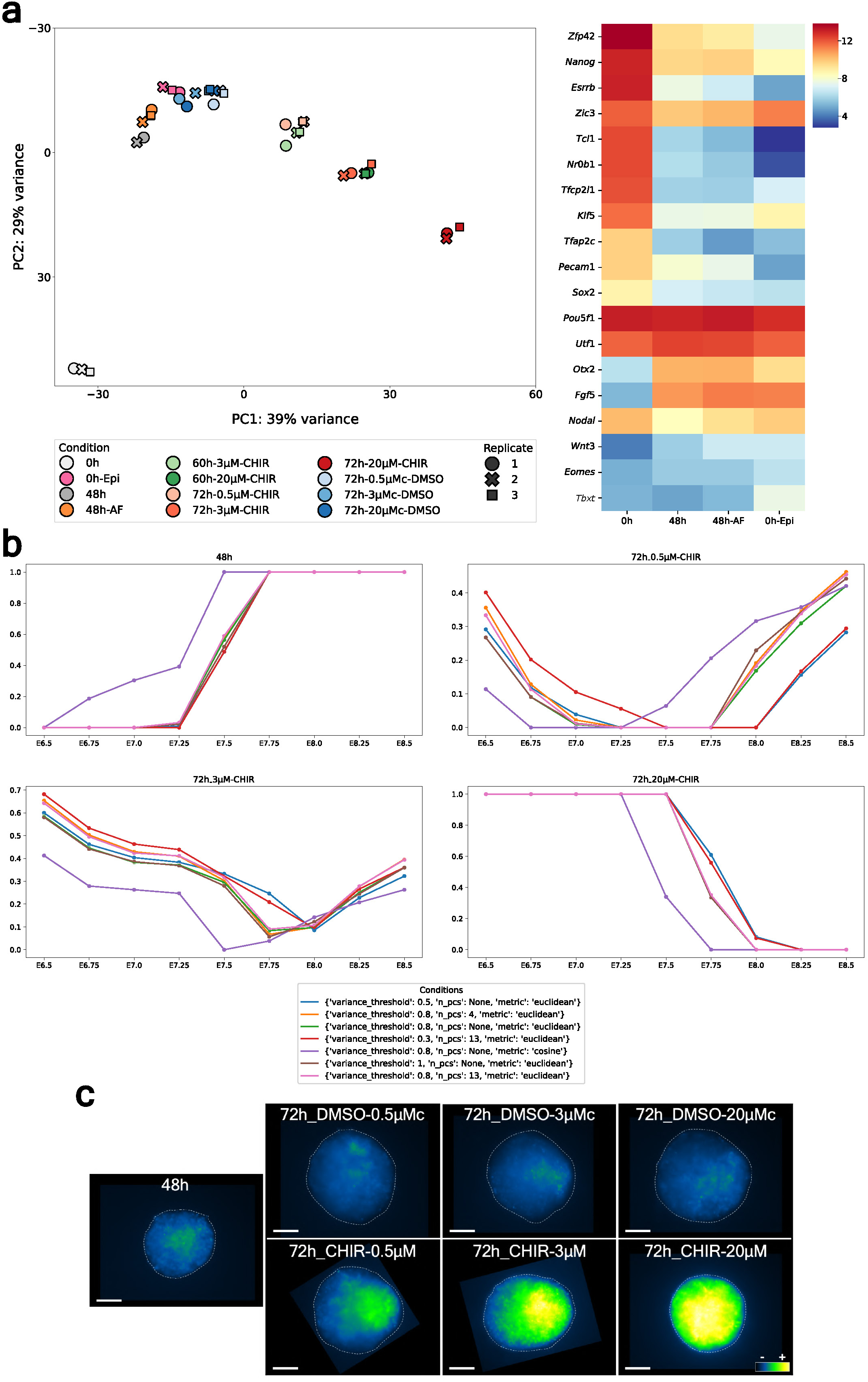
Deconstructing and modulating the mouse gastruloid model system. **a**, Transition from naive to primed pluripotency occurs between 0 and 48h of the gastruloid protocol. PCA of gastruloid and EpiLSCs (Epi) bulk RNA sequencing samples (three replicates per condition, from three independent experiments). mESCs at the beginning of the gastruloid protocol (0h) form a distinct cluster, far away from all the other samples. EpiLSCs cluster between the 48 and 72h DMSO gastruloids, suggesting that a transition from naive to formative/primed pluripotency occurs between 0 and 48h in the standard gastruloid protocol. The addition of ActA and Fgf (AF; see **Methods**) during this time window (0-48h) does not result in significant transcriptomic differences as these gastruloids at 48h cluster between the 48h control gastruloids and the EpiLSCs samples. A gene expression heatmap (mean of the replicates variance-stabilised read counts) illustrating different states of pluripotency is also provided. Contrary to all 48h gastruloids and EpiLSCs samples, mESCs (0h) display high expression of genes associated with naive pluripotency (e.g. *Zfp42*, *Nanog* and *Essrb*) and low levels of epiblast-related genes (e.g. *Otx2* and *Fgf5*). EpiLSCs exhibit slightly higher expression of PS-related genes (e.g. *Tbxt*) compared to 48h gastruloids, further distinguishing their transcriptomic profile. **b**, Robustness analysis of gastruloids and embryos transcriptomic distance. Correlation between pseudo-bulk data from Pijuan-Sala et al.^86^ and 48h gastruloids,72h gastruloids treated with 0.5, 3 and 20µM CHIR. Different choices of three analysis parameters were also tested: 1) variance threshold, which selected the number of genes to use for the PCA (Fig. 1d), 2) number of PCA components and 3) metric to compute the pairwise distance between the embryo and gastruloid samples. All the parameter choices lead to similar results, with 48h gastruloids being closer to early-stage embryos and 72h gastruloids close to late-stage embryos, in a CHIR concentration manner. The main outlier is the parameter set using a different metric (cosine metric), which displays a similar trend but shifts the expected minimum at each CHIR concentration to earlier stages in the 0.5 and 3µM CHIR samples. **c**, Wnt signalling activity in CHIR and DMSO-treated gastruloids. Representative maximum intensity projections (MIPs) of TCF/Lef reporter gastruloids at 48h and 72h (total of 60 gastruloids imaged per condition, from three independent experiments), upon different CHIR and DMSO treatments provided at 48h. Contrary to DMSO controls (c), which do not show a significant change in Wnt activity, CHIR-treated gastruloids display higher reporter expression in a concentration-dependent manner. The scale bar is equivalent to 100µm.

**Extended Data Fig 2.**
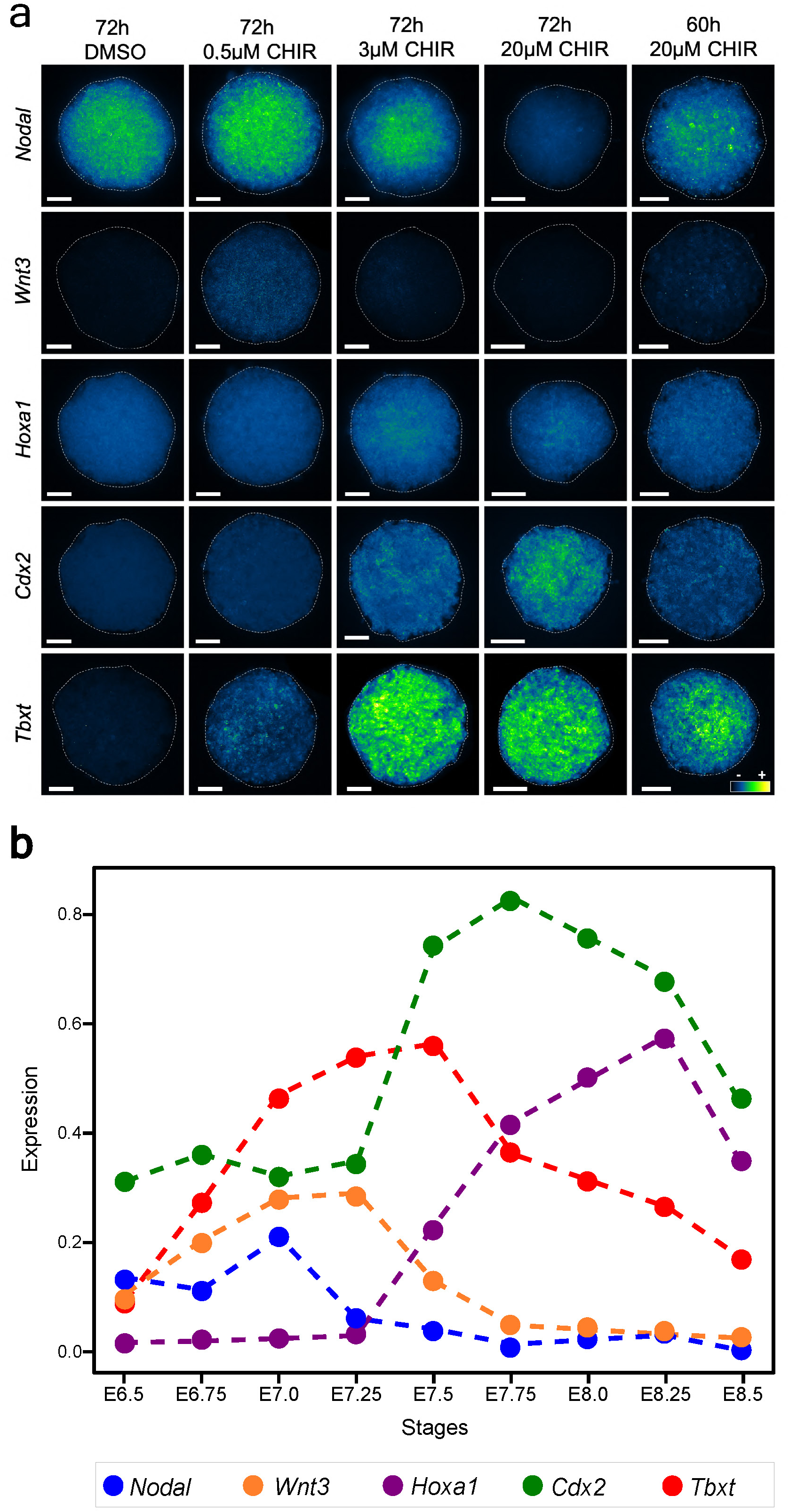
Expression of gastrulation marker genes in mouse embryos and gastruloids. **a,** HCR (MIP) for early and late PS-related genes in gastruloids made with different levels of Wnt signalling. At 72h, gastruloids treated with DMSO do not express any of the late PS marker genes (e.g. *Cdx2*), with only *Nodal* being detected. In contrast, CHIR gastruloids show an upregulation of *Tbxt* and *Cdx2* and a downregulation of genes associated with early developmental stages (e.g. *Nodal* and *Wnt3*) in a concentration-dependent manner. These changes promoted by Wnt signalling seem to occur also in a temporal manner as 20µM CHIR-treated gastruloids at 60h display a gene expression resembling that of 3µM CHIR-treated gastruloids at 72h (e.g. low *Nodal* and *Cdx2,* high *Tbxt*). The MIPs shown here are representative images from two independent experiments, with 4 gastruloids imaged per condition/experiment. The scale bar is equivalent to 50µm. **b**, Pseudo-bulk temporal expression of specific gastrulation-related genes in mouse embryos (containing extra-embryonic tissues). The expression of the early PS marker genes *Nodal* and *Wnt3* significantly decreases after E7.0, with the upregulation of *Tbxt*, *Cdx2* and *Hoxa1*. The drop in the levels of *Tbxt* after E7.5 is consistent with the regression of the PS. High levels of *Cdx2* at early stages are consistent with its expression in extra-embryonic tissues. The gene expression values displayed in the figure were derived from a scRNA-seq mouse atlas^86^ and are the mean expression calculated over all the cells assigned to each timepoint, with the data normalised and logp1-fold scaled. Dashed lines between timepoints represent linear interpolations to aid visualisation.

**Extended Data Fig 3.**
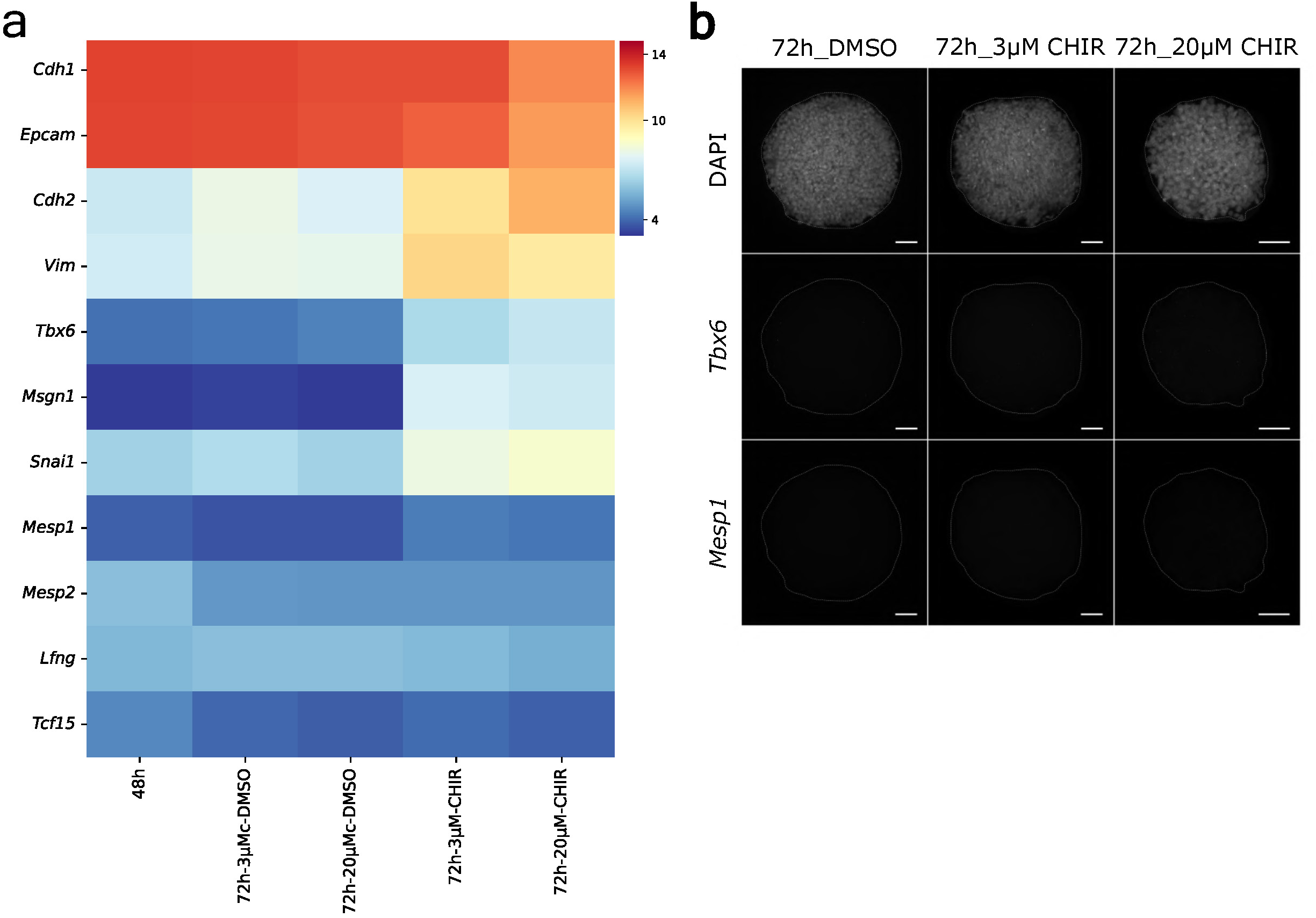
Wnt signalling is not primarily involved in mesoderm specification. **a**, Gene expression heatmap (mean variance-stabilised read counts of replicates) of gastruloid bulk RNAseq samples at 48h and 72h (three replicates per condition, from three independent experiments). Mesenchymal markers, such as *Cdh2*, *Vim*, and *Snai1*, are upregulated upon CHIR treatment. Notably, *Cdh2* expression increases in a concentration-dependent manner, with the highest expression observed with 20μM CHIR. In contrast, the expression of epithelial-related genes (*Cdh1* and *Epcam*) decreased when CHIR was used. Despite this classical EMT profile, the levels of key mesodermal-related genes remain low (e.g. *Tbx6*, *Msgn1, Mesp1* and *Tcf15)*, even when at higher concentrations of CHIR. **b**, HCR (MIP) showing the lack of significant *Mesp1* and *Tbx6* expression in 72h gastruloids, including those treated with 20μM CHIR. The MIPs shown here are representative images of two independent experiments, with 3-4 gastruloids imaged per condition/experiment. The scale bar is equivalent to 50µm.

**Extended Data Fig 4.**
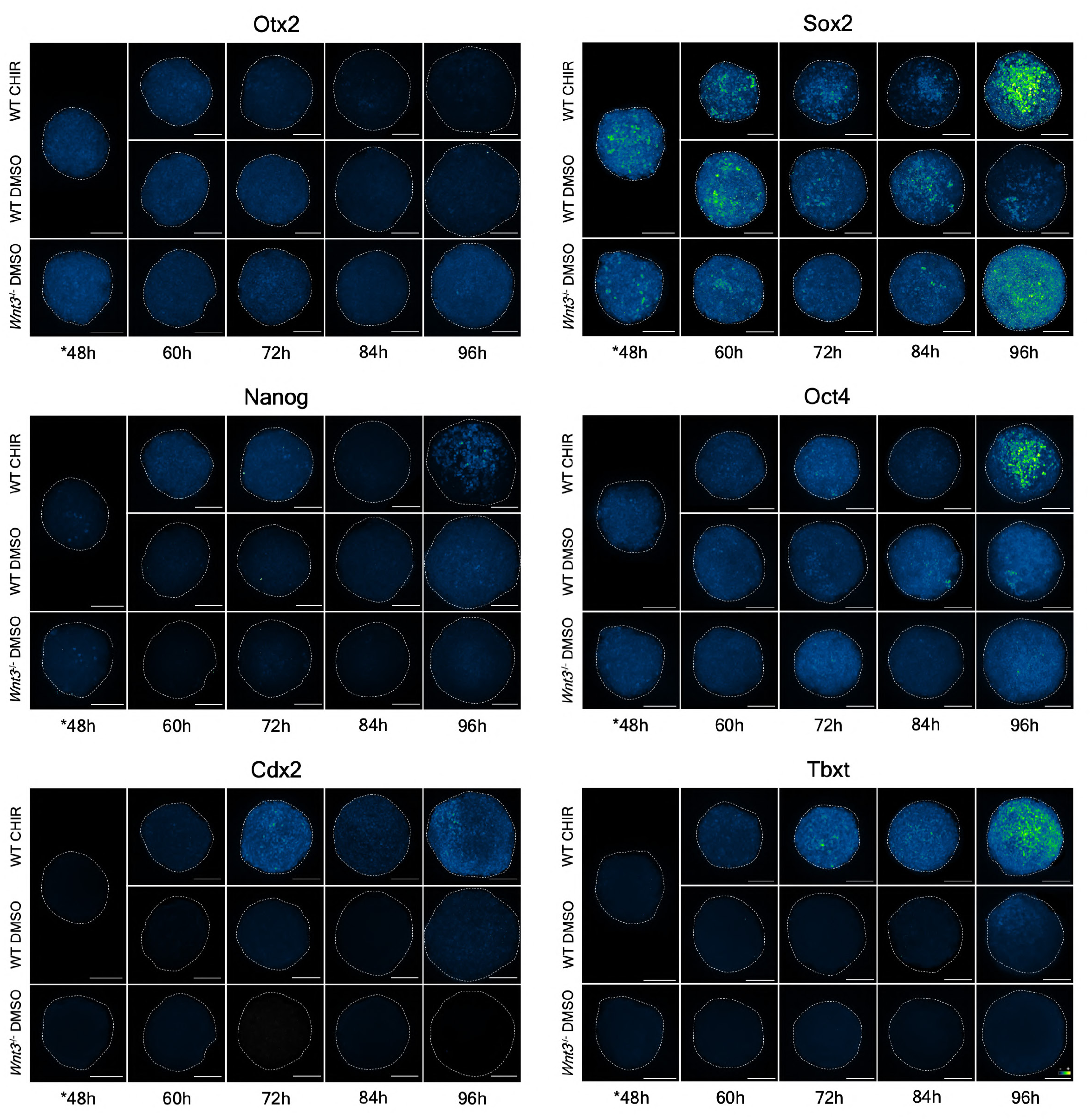
Assessing the role of Wnt signalling in the maturation of mouse pluripotent epiblast cells in gastruloids. Immunofluorescence stainings (MIP) for Nanog, Oct4, Cdx2 and Tbxt, Sox2 and Otx2, performed in WT-CHIR, WT-DMSO and Wnt3KO-DMSO treated gastruloids (see **Methods** for details). WT CHIR/DMSO gastruloids at 48h (*48h) are similar because the treatments were only given between 48h and 72h. At this stage, WT and *Wnt3* mutant gastruloids display similar expression patterns for all tested markers. Otx2, Oct4 and Sox2 are broadly expressed, though Sox2 levels are higher in a subset of cells. Nanog is weakly expressed, with only a few cells showing elevated levels. Tbxt and Cdx2 are not expressed. At 60h, the expression patterns remain largely similar across conditions, but some differences begin to emerge. The levels of Cdx2, Tbxt and Nanog were slightly higher in WT gastruloids treated with CHIR and their expression, especially that of Tbxt and Cdx2, continues to increase over time in these gastruloids. In contrast, there is a concomitant downregulation of Otx2, starting at 72h. WT-DMSO gastruloids exhibit a similar pattern later in time: the Otx2 reduction starts only at 84h, and the levels of Nanog, Tbxt and Cdx2 start to come up only at 96h. In contrast, *Wnt3* mutant gastruloids maintain the expression of Otx2 and Oct4 throughout the entire protocol, display very low levels of Nanog and do not show any expression of Cdx2 and Tbxt. Also, the expression of Sox2 sees a significant increase at 96h. Regarding WT standard gastruloids, the expression of Oct4 and Sox2 decreases after the addition of CHIR, being only positive in some cells at 84h. WT-DMSO gastruloids exhibited a similar Sox2 phenotype but only at 96h. At this time, WT-CHIR gastruloids display high levels of Sox2 in specific groups of cells, where it is also possible to observe high expression of Tbxt, Oct4 and Nanog, which resembles the emergence of neuromesodermal progenitors from the pluripotent epiblast. The scale bars represent 100µm.

**Extended Data Fig. 5.**
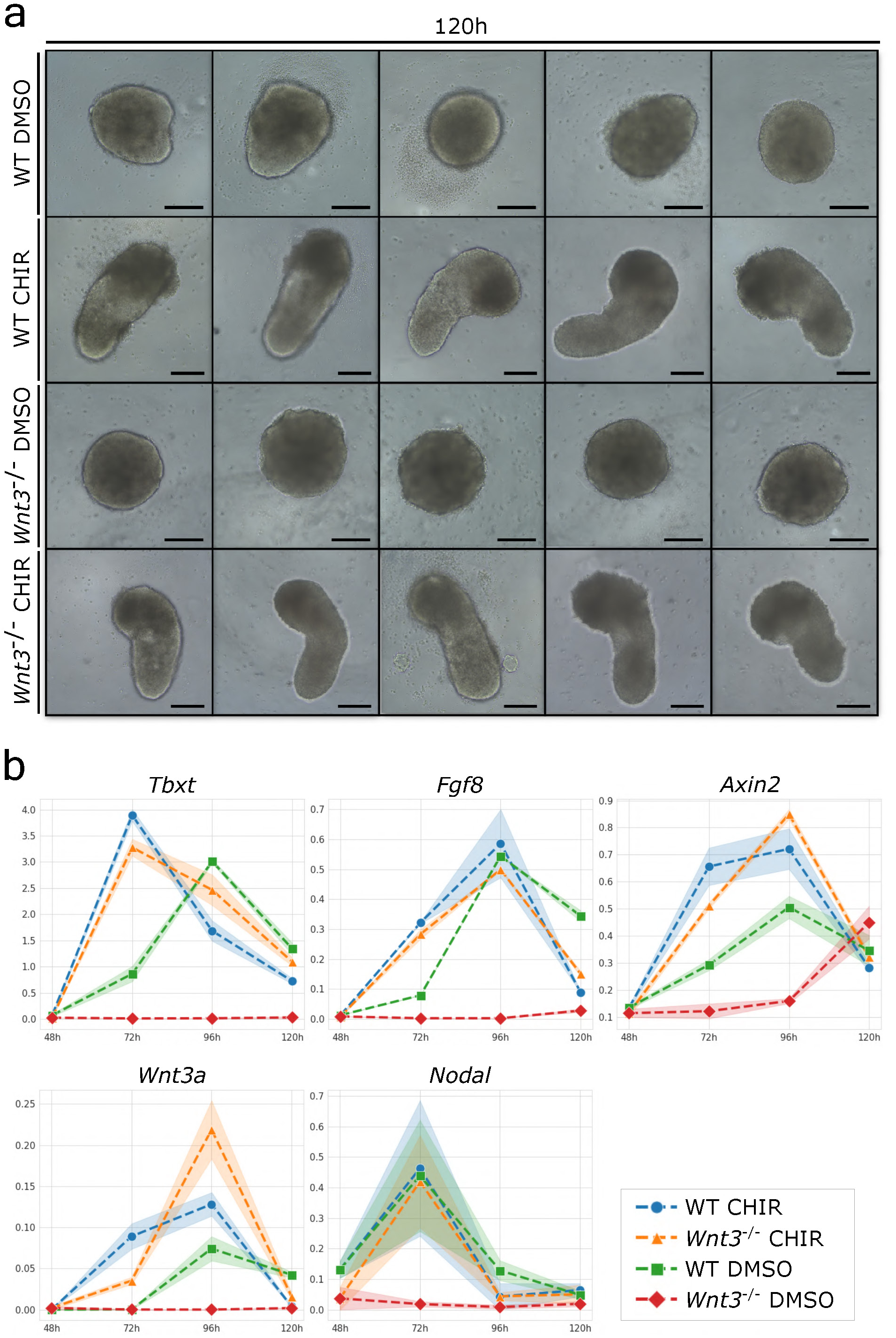
Characterisation of *Wnt3* mutant mouse gastruloids. **a**, Representative images from 5 independent experiments (total of around 250 gastruloids per condition) highlighting distinct morphological differences between WT and *Wnt3*^-/-^ gastruloids at 120h, treated with either 3µM CHIR or a corresponding amount of DMSO (48-72h). The scale bars represent 200µm. **b**, Line plots showing the temporal expression profiles of *Tbxt*, *Fgf8*, *Axin2*, *Wnt3a* and *Nodal* via qPCR in WT and *Wnt3^-/-^* gastruloids treated with either DMSO or CHIR (48-72h) at 48h, 72h, 96h and 120h (three replicates per experiment, two independent experiments). *Tbxt* expression peaks at 72h in both WT and *Wnt3 mutant* gastruloids treated with CHIR, and at 96h in WT-DMSO gastruloids. *Wnt3*^-/-^ gastruloids do not exhibit any significant levels of *Tbxt* throughout the gastruloid protocol. Similarly, the expression of *Fgf8*, *Wnt3a* and *Nodal* is highly limited in *Wnt3*KO-DMSO gastruloids. *Wnt3* mutant gastruloids treated with CHIR display expression patterns similar to WT gastruloids, particularly those treated with CHIR. *Axin2* is upregulated between 96h and 120h in *Wnt3* mutant gastruloids treated with DMSO, reaching levels comparable to other conditions by 120h. Log1p-fold normalisation was used on the y-axis. Shaded regions in the plots represent error margins, and dashed lines between timepoints indicate linear interpolations to aid visualisation.

**Extended Data Fig. 6.**
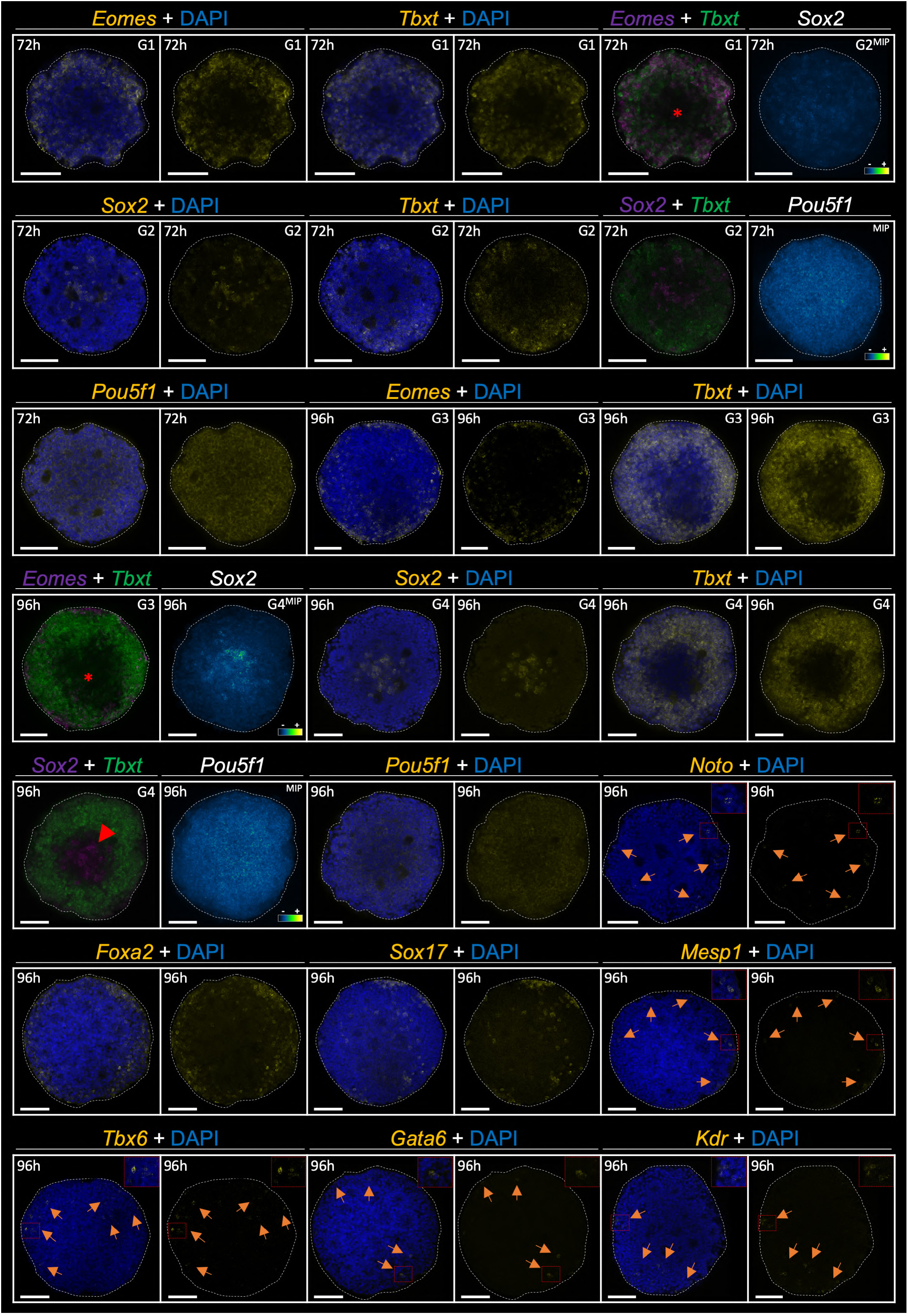
Radial cell fate specification in Activin A treated gastruloids. Representative HCR images from medial optical sections unless stated otherwise (MIP) from three independent experiments, with around 3-4 gastruloids imaged per condition/experiment. At 72 hours, ActA induces a radial organisation in gastruloids as indicated by the expression of *Eomes* and *Tbxt* across most of the gastruloid except for its central region, which contains some *Sox2*-positive cells. *Oct4* is uniformly expressed throughout the gastruloid at this stage. The radial asymmetry becomes more pronounced by 96h, with a distinct core of *Sox2*-positive cells emerging in the centre, surrounded by *Tbxt*-expressing cells. *Eomes* expression is predominantly localised in the outer regions, while *Oct4* continues to be expressed throughout the gastruloid. At this stage, derivatives of the early PS are observed in the periphery of the ActA-treated gastruloids. These include *Noto* and *Foxa2*, marking node/notochord-like cells, *Foxa2* and *Sox17* for endodermal lineage, *Kdr* for haematoendothelial progenitors, and *Tbx6* and *Mesp1* for anterior mesoderm. Inlets provide magnified views of these expression patterns. "G" denotes gastruloid. The scale bar represents 50 µm.

**Extended Data Fig 7.**
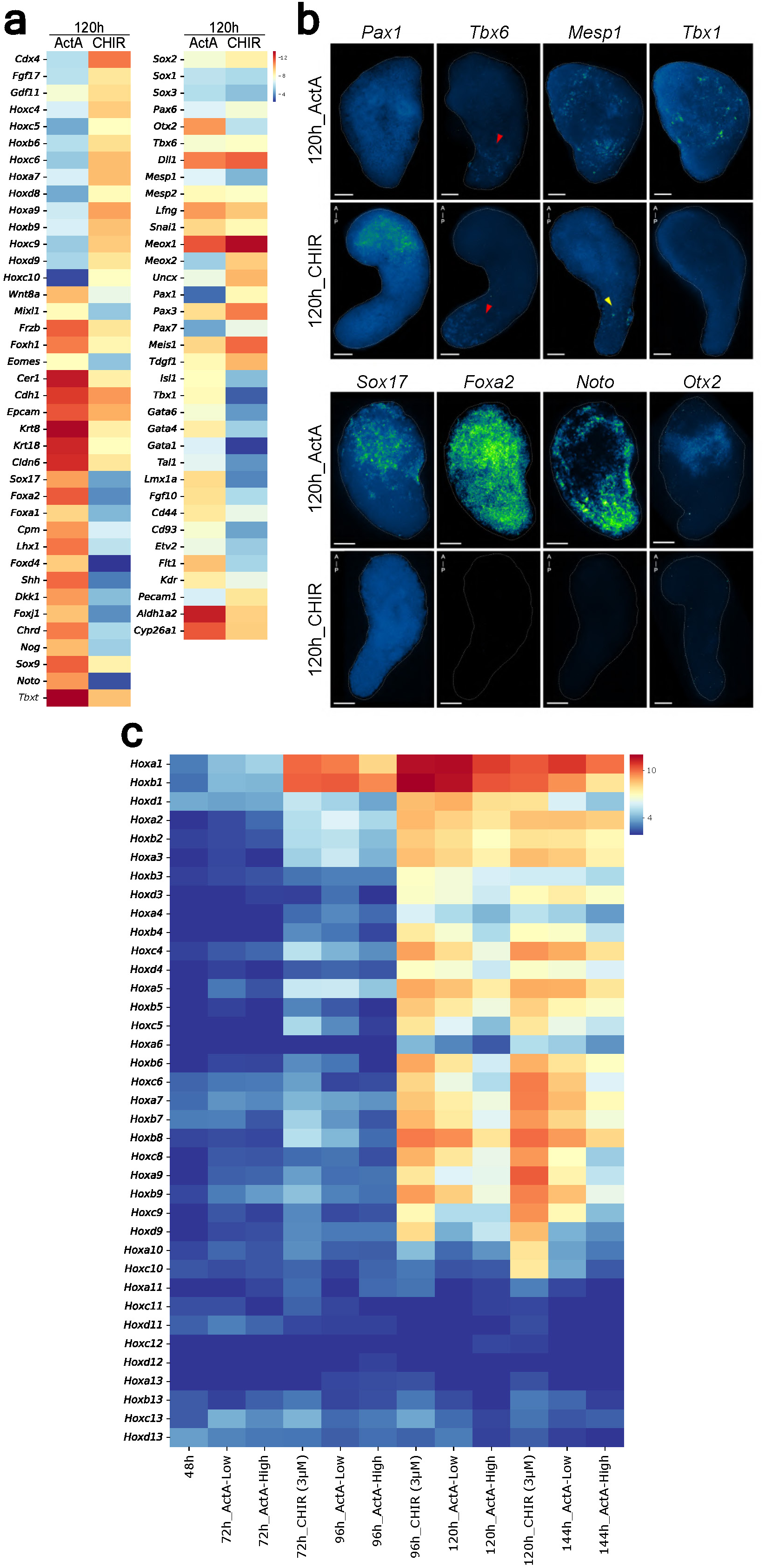
Gene expression comparison between Activin A and CHIR-treated gastruloids. **a**, Heatmap showing gene expression (mean of variance-stabilized read counts across replicates) from bulk RNAseq of 120h gastruloids treated with (3 µM) CHIR or (high) ActA (three replicates per condition, from three independent experiments). Unlike ActA, the CHIR treatment leads to a gene expression signature characteristic of derivatives of the late PS. For example, genes associated with the caudal epiblast, such as *Cdx4*, *Fgf17*, and *Gdf11*, are exclusively expressed in CHIR-treated gastruloids. Additionally, *Hoxc10 –* a tailbud marker - is expressed only in gastruloids made with CHIR, suggesting they are more developmentally advanced and posteriorized compared to ActA-treated gastruloids. The expression of several Hox genes, including *Hoxc4*, *Hoxb6*, *Hoxd8*, and *Hoxa9*, in CHIR-treated gastruloids further supports this posteriorization. Conversely, the presence of *Wnt8a*, a late gastrulation marker, in ActA but not CHIR-treated gastruloids suggests a temporal delay in the first ones. Moreover, the expression of *Sox2*/*Pax6*, and *Uncx*/*Meox1*/*Lfng*/*Tbx6* in CHIR-treated gastruloids indicates the development of trunk neural and mesodermal-like cells, respectively. In contrast, ActA-treated gastruloids lack *Pax6* expression, which suggests an absence of trunk neural tissue. The mesodermal gene expression signature in these gastruloids includes *Tbx1*, indicative of its anterior identity. The expression of *Sox17*, *Foxa2*, *Noto*, *Gsc* and *Chrd* further emphasises the anteriorization of ActA-treated gastruloids compared to those treated with CHIR. Additionally, *Kdr* and *Flt1* expression in ActA-treated gastruloids suggests the presence of endothelial-like cells. **b,** HCR (MIP) images illustrate gene expression in 120h gastruloids treated with ActA or CHIR. *Pax1* is expressed in the anterior compartment of CHIR-treated gastruloids but absent in ActA gastruloids. *Tbx6* is expressed in the posterior regions of both gastruloid types (indicated by red arrowheads), indicating mesoderm formation. *Mesp1* is expressed in the determination front of gastruloids made with CHIR (yellow arrowhead) but shows a distinct pattern in ActA-treated gastruloids, next to *Tbx1* expression. *Sox17*, *Foxa2* and *Noto* expression in ActA-treated gastruloids highlights the spatial distribution of endoderm and node/notochord-like cells, respectively. Expression of *Otx2* exclusively in gastruloids made with ActA is also in accordance with the bulk RNAseq data. The MIPs shown here are representative images of three independent experiments, with 3 gastruloids imaged per condition/experiment. The scale bar is equivalent to 50µm. **c**, Comparative temporal *Hox* gene expression in CHIR and ActA-treated gastruloids. No *Hox* gene expression is detected at 48h. By 72h, *Hoxa1* and *Hoxb1* begin to be expressed in CHIR-treated gastruloids but not in gastruloids made with ActA, even when a high dose is used. At 96 hours, ActA-treated gastruloids begin expressing *Hoxa1* and *Hoxb1*, though expression levels vary inversely with the dose of Activin. CHIR-treated gastruloids exhibit at this stage a broader *Hox* gene profile, including the expression of some posterior ones like *Hoxb9*. At 120 hours, CHIR-treated gastruloids show a highly posteriorized *Hox* gene signature, with significant levels of *Hoxc10 (*tailbud marker). In contrast, gastruloids treated with a low dose of ActA express a *Hox* gene signature similar to that of 96h CHIR-treated gastruloids, with notable expression of posterior *Hox* genes like *Hoxa5* and *Hoxb9*, but not *Hoxc5*, *Hoxc6*, or *Hoxa/c/d9*. Gastruloids exposed to a high dose of ActA exhibit a more limited *Hox* gene expression, predominantly featuring anterior *Hox* genes such as *Hoxb1*, *Hoxa3* and *Hoxa5*. By 144 hours, gastruloids made with a low dose of ActA show expression of several posterior *Hox* genes, though not *Hoxc10*. Meanwhile, gastruloids treated with high levels of ActA continues to display a strong anterior *Hox* gene signature, with low levels of posterior *Hox* genes.

**Extended Data Fig. 8.**
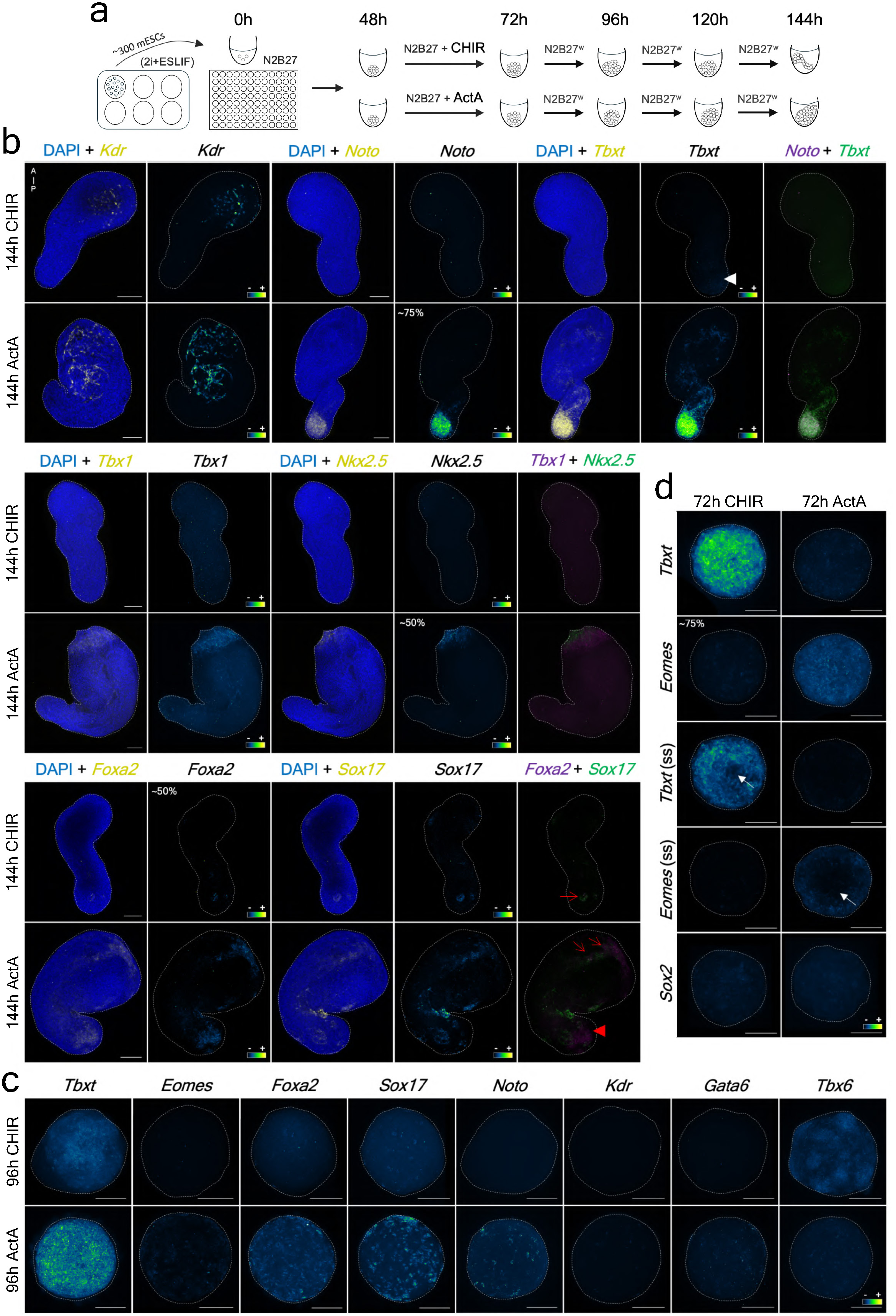
Robustness of CHIR and Activin A treated gastruloids. **a**, Scheme highlighting the culturing conditions. 129/svev mESCs were cultured in 2i+ESLIF, and CHIR and ActA gastruloids were developed using homemade N2B27 media (see **Methods**). **b**, HCR representative MIP images (between 7-9 gastruloids imaged per condition, from three independent experiments) illustrating the differences of the above-mentioned gastruloids at 144h, treated with CHIR (standard) or High ActA between 48h and 72h. *Kdr* was expressed in both types of gastruloids, though more in gastruloids treated with ActA. The expression of *Foxa2* and *Sox17* was low and variable in CHIR-treated gastruloids, with only around 50% of the gastruloids displaying significant levels of *Foxa2*. Co-localisation of *Sox17* and *Foxa2* (in white/greyish, red arrow) is suggestive of definitive endoderm-like cells, and is significantly enhanced in gastruloids treated with ActA. The expression of *Foxa2* in regions of ActA gastruloids lacking *Sox17* (red arrowhead) is suggestive of notochord-like cells. This is supported by the expression of *Noto*, which was only detected in 75% of the imaged ActA gastruloids. Similarly to Fig. 4 and 5, the expression of *Tbxt* is also observed in more anterior regions of the gastruloids treated with ActA that lack the expression of *Noto*. In contrast, *Tbxt* is only expressed at low levels in the posterior part of CHIR gastruloids (white arrowhead). The expression of the anterior cardiac markers *Tbx1* and *Nkx2.5* was only noticed in ActA-treated gastruloids, with *Nkx2.5* present in around 50% of the imaged gastruloids. Morphologically, ActA gastruloids exhibited a significant expansion in the anterior compartment compared to those treated with CHIR. The scale bars represent 100µm. **c**, HCR representative MIP images (between 6-8 gastruloids imaged per condition, from three independent experiments) of ActA and CHIR gastruloids at 96h. Similarly to Fig. 3, ActA gastruloids display a stronger expression of gene markers for cell fates known to arise from the early PS than those treated with CHIR (e.g. *Foxa2*, *Sox17* and *Noto*). Gastruloids treated with CHIR express higher levels of *Tbx6* and a polarised *Tbxt* expression. The scale bars represent 100µm. **d**, Analysis of *Eomes* and *Tbxt* expression in 72h CHIR and ActA gastruloids via HCR; representative MIP images (except when “MOS” is indicated – media optical section) of 8 gastruloids imaged per condition, from three independent experiments. The levels of *Tbxt* are higher in gastruloids treated with CHIR, whereas the expression of *Eomes* is enhanced in ActA gastruloids. The results are similar to those highlighted in Fig. 3f, though around 75% of the imaged 129/svev CHIR gastruloids display significant levels of *Eomes* at 72h, which likely explains the appearance of some anterior cell fates in these gastruloids. Evidence for radial asymmetry (see also **Extended Data Fig. 6**) in the expression of *Tbxt* and *Eomes* can be noticed in both types of gastruloids (white arrow), though it is more enhanced in those treated with ActA. The scale bars represent 100µm.

**Extended Data Fig. 9.**
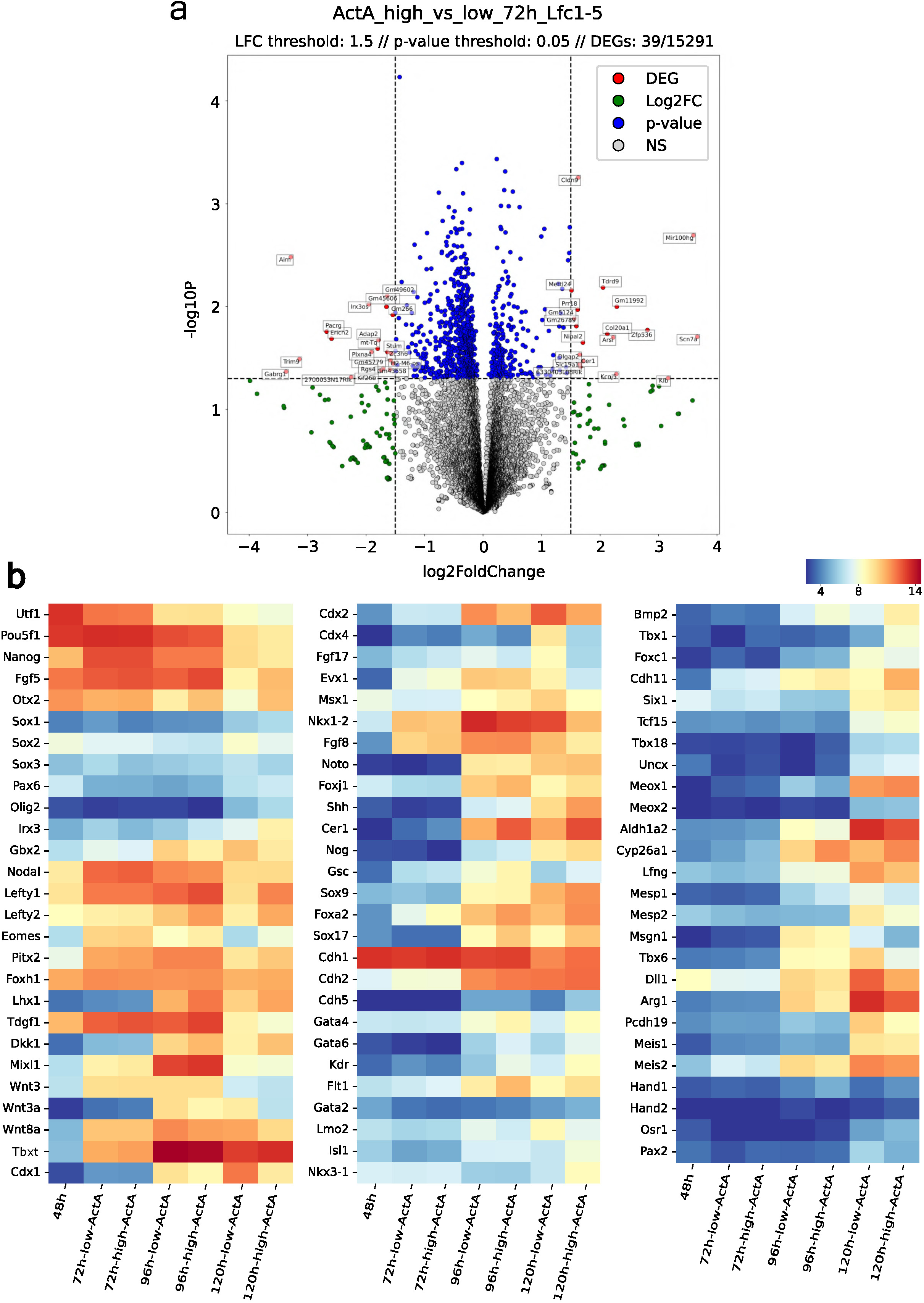
Comparative gene expression analysis of gastruloids treated with low or high levels of Activin A. **A**– Volcano plot of ActA (ActA) gastruloids bulk RNAseq data (three replicates per condition, from three independent experiments) highlighting the small difference between gastruloids treated with high versus low levels of ActA at 72 hours. Red dots represent differentially expressed genes (DEG) meeting the criteria of log fold change (LFC) greater than 1.5 and a p-value below 0.05; non-significant genes (NS; do not meet the previous conditions) are shown in grey. Out of 15,291 genes analysed, only 39 were considered significant based on these thresholds. **B** – Heatmap illustrating gene expression (mean of variance-stabilised read counts across replicates) from bulk RNAseq of 48h gastruloids and 72-144h gastruloids treated with either low or high levels of ActA. At 72h, a slight upregulation of *Otx2*, *Nodal*, *Tdgf1*, *Tbxt*, *Fgf8*, *Evx1*, *Eomes*, and *Pitx2* expression is observed in gastruloids treated with a higher amount ActA, with the most notable increase seen in the expression *Foxa2*. By 96h, the differences become more pronounced, with high ActA-treated gastruloids showing elevated expression of *Otx2*, *Lefty1/2*, *Tdgf1*, *Eomes*, *Lhx1*, *Dkk1*, *Msx1*, *Cer1*, *Gsc*, *Foxa2*, *Sox17* and *Gata4*, indicative of a more anterior patterning. In contrast, gastruloids treated with low ActA upregulate genes associated with the late PS, like *Wnt3a*, *Cdx1*, *Cdx2* and *Meis2*. Of note, ActA gastruloids express higher levels of *Cyp26a1*, which is involved in the degradation of retinoic acid (RA). At 120h, the divergence between the two conditions becomes even more distinct. Gastruloids exposed to higher levels of Nodal signalling continue to express higher levels of genes associated with the development of anterior body structures like *Dkk1* and *Lhx1*, as well as endothelial-related genes like *Kdr* and *Flt1*. The higher levels of *Foxa2*, *Sox17*, *Shh* and *Nog* expression in gastruloids developed with a high dose of ActA suggests the presence of more endoderm and notochord-like cells in these gastruloids. Conversely, low ActA-treated gastruloids show increased expression of *Wnt3a*, *Wnt8a*, *Cdx1/2/4*, *Fgf17*, and *Nkx1-2*, markers linked to the late PS. Gastruloids treated with higher ActA continue to maintain significantly higher levels of *Cyp26a1*, suggesting greater protection against RA activity in comparison to those treated with a lower ActA dose. High levels of *Lefty1* in 120h gastruloids made with a high ActA suggest again that Nodal signalling activity might be more elevated in comparison to low ActA-treated gastruloids.

**Extended Data Fig 10.**
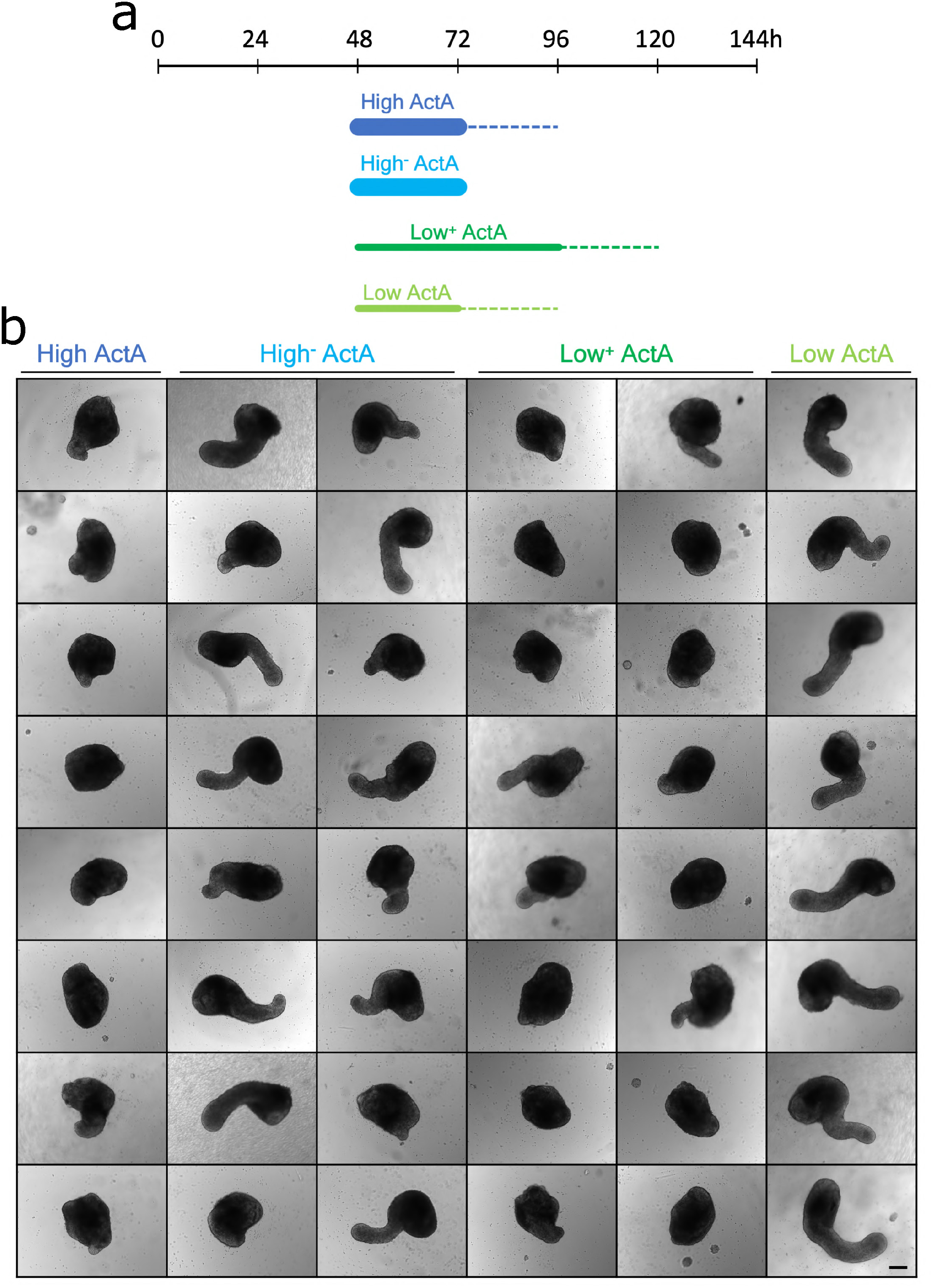
Temporal and dose-dependent effects of Activin A in mouse gastruloids. **a**, Schematic diagram of different ActA-treated gastruloids. In contrast with High ActA gastruloids, which were developed according to the standard protocol (see **Methods**), High^-^ ActA-treated gastruloids were washed 2 times at 72h with fresh N2B27 media. These washes were designed to remove any traces of ActA from the media, thus limiting its effect in time in comparison with standard High ActA gastruloids. To further test the temporal effects of ActA in mouse gastruloids, this recombinant protein was also introduced at lower concentrations (similar to Low ActA-treated gastruloids) between 72h and 96h – Low^+^ ActA gastruloids. **b**, Representative images of High, High^-^, Low^+^ and Low ActA-treated gastruloids at 144h (three independent experiments, 40 gastruloids imaged per condition). In contrast to Low ActA gastruloids, those treated with high levels of ActA tend to display mostly a round shape and exhibit minimal elongation at the posterior part. The formers display an enhanced elongated phenotype. Reducing the temporal effect of high levels of Nodal signalling at 72h has a significant effect because High^-^ ActA gastruloids display a phenotype more similar to Low ActA-treated gastruloids (more predominant posterior elongations), in comparison with High ActA gastruloids. Similarly, increasing the time of the ActA treatment up to 96h also affects the phenotype of gastruloids at 144h. Low+ ActA gastruloids exhibit a phenotype that resembles that of High ActA-treated gastruloids. The scale bar is equivalent to 50µm.

**Supplementary Data Figure 1.**
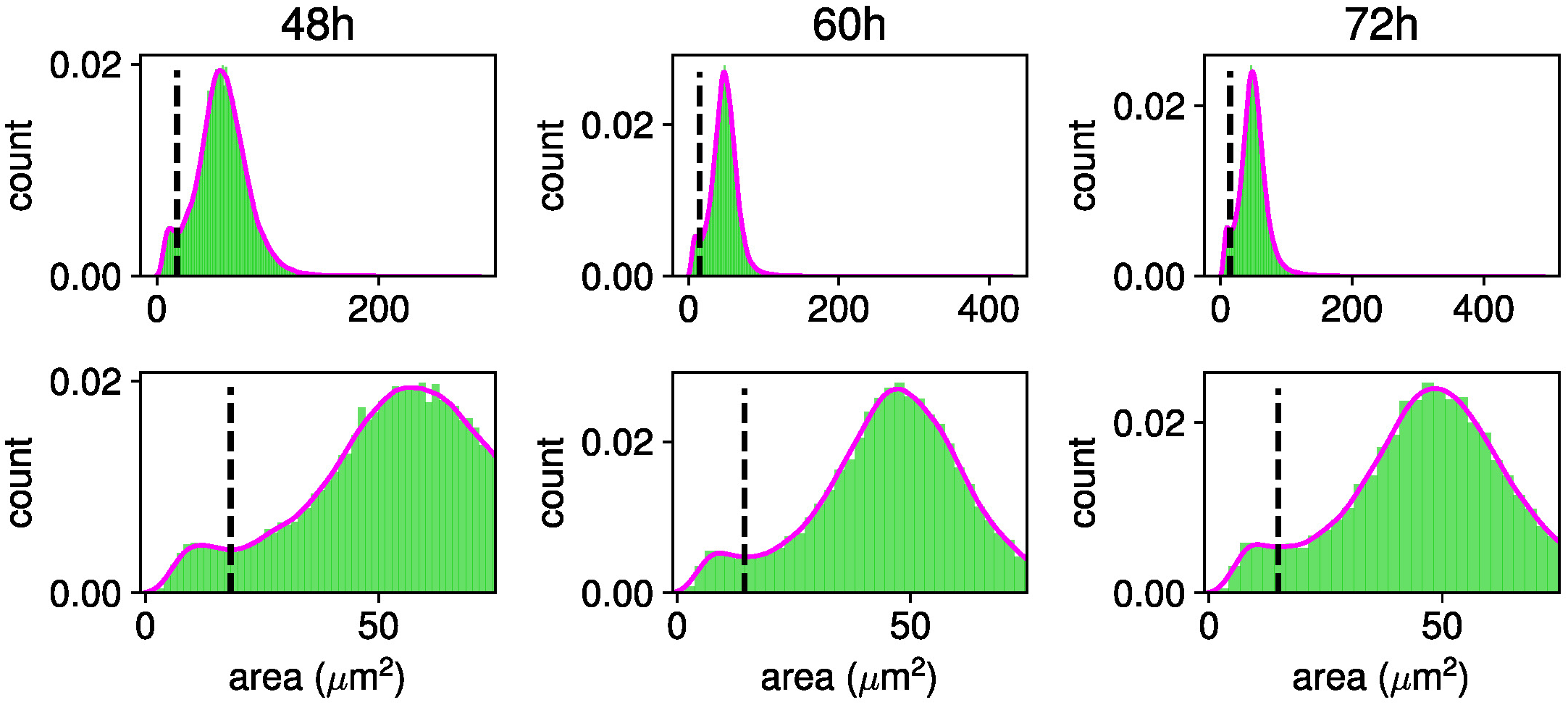
Debris removal during 3D image analysis. Distributions of segmented object areas from all gastruloids along the x-axis at the different experimental time points. The histograms at the top show the area distributions, overlaid with a KDE using the smallest bandwidth that results in a single local minimum. This minimum, marked by a vertical dashed line, indicates the threshold separating cellular debris from intact cells. A zoom over the x-axis is provided below to improve the visualisation of the existing bimodality.

**Supplementary Data Figure 2.**
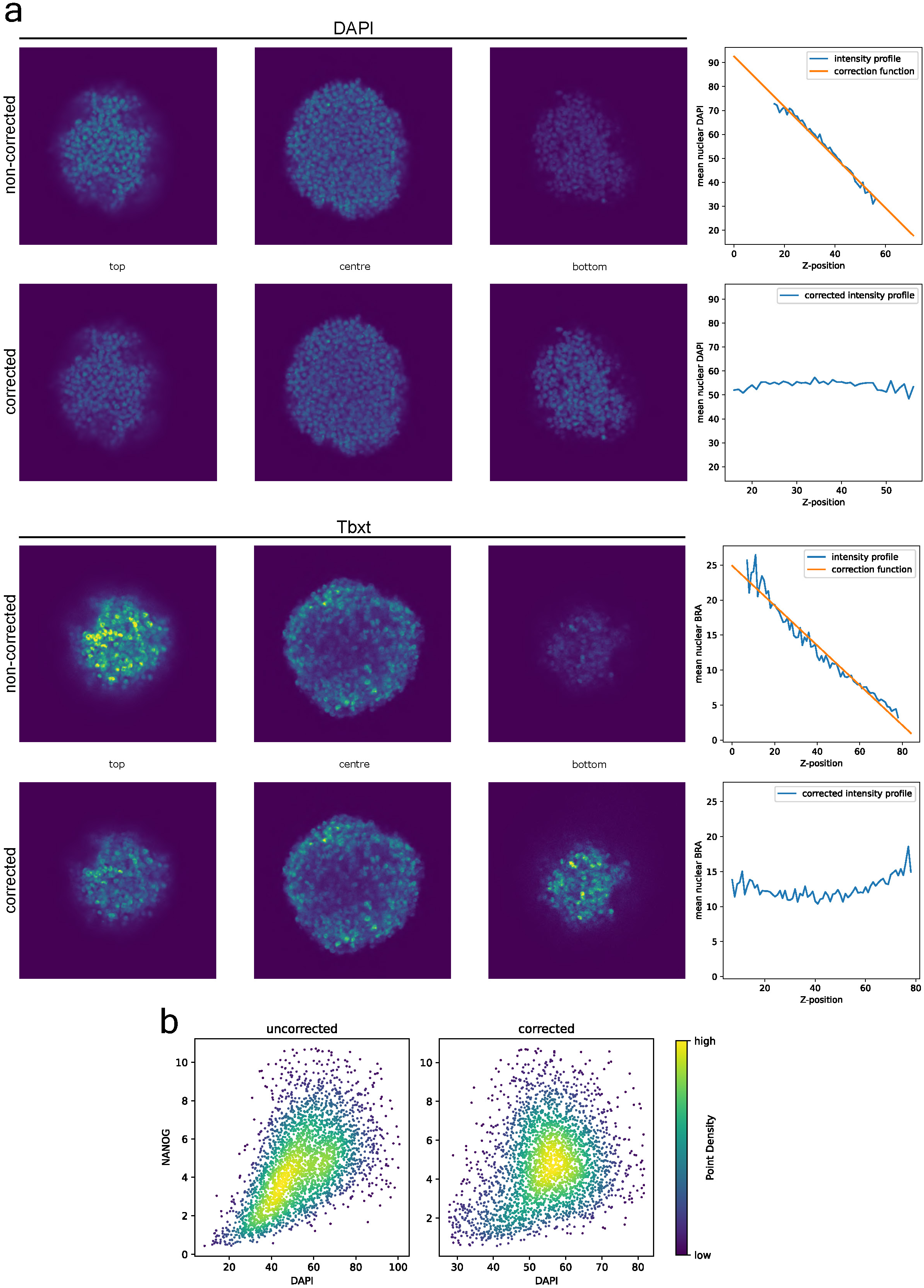
Signal correction in 3D gastruloid datasets. **a**, Example of immunofluorescence optical sections of the DAPI (48h WT gastruloids) and the Tbxt antibody channel (72h WT CHIR-treated gastruloid) before and after signal drift correction along the z-axis. Quantification plots show the mean nuclear expression along the z-axis, before and after signal correction. **b**, Scatter plot displaying the expression of Nanog and DAPI (48h WT gastruloid) with and without the signal drift correction across the z-axis.

**Supplementary Data Figure 3.**
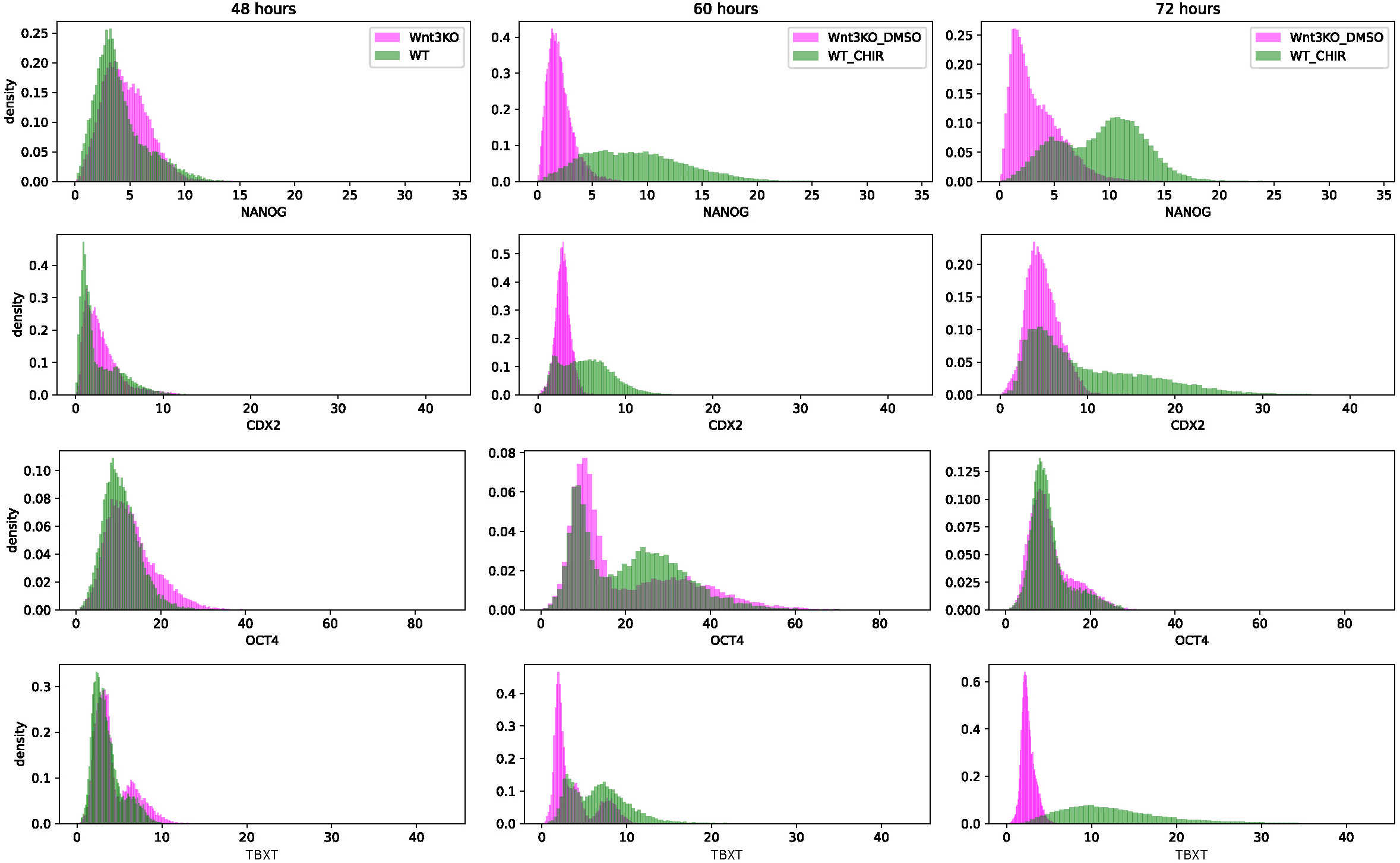
Intensity histograms of 3D segmented gastruloids after signal correction. The histograms shown here are similar to those in Fig. 2a, but the intensity values were adjusted following signal drift correction across the z-axis (see **Methods**).

**Supplementary Data Figure 4.**
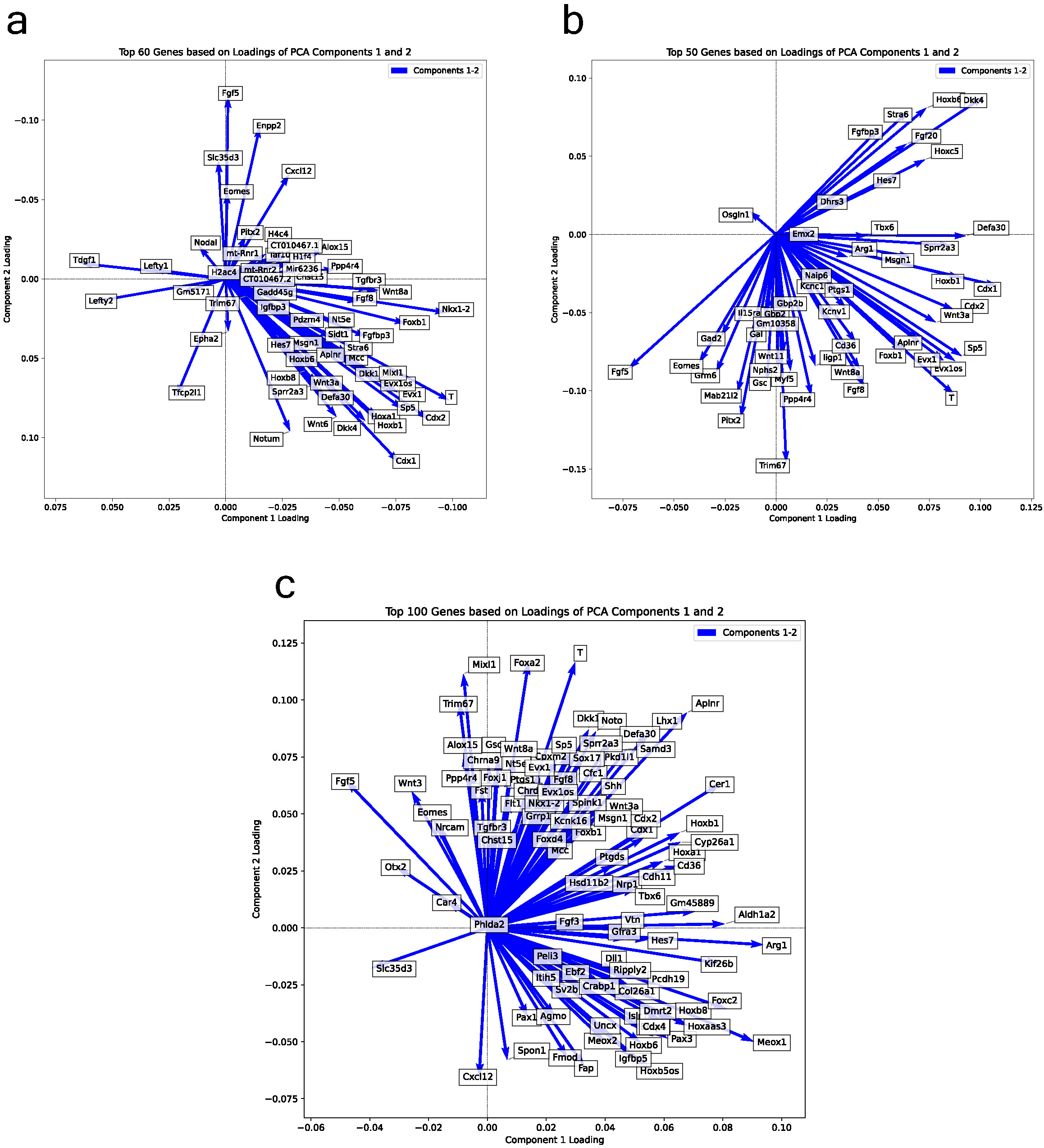
Full top gene loadings derived from the various principal components. **a**, Top 60 gene loadings related to the PCA shown in Fig. 1b. **b**, Top 50 gene loadings related to the PCA shown in Fig. 1d. **c**, Top 100 gene loadings related to the PCA shown in Fig. 3b.

**Supplementary Data Table 1.**
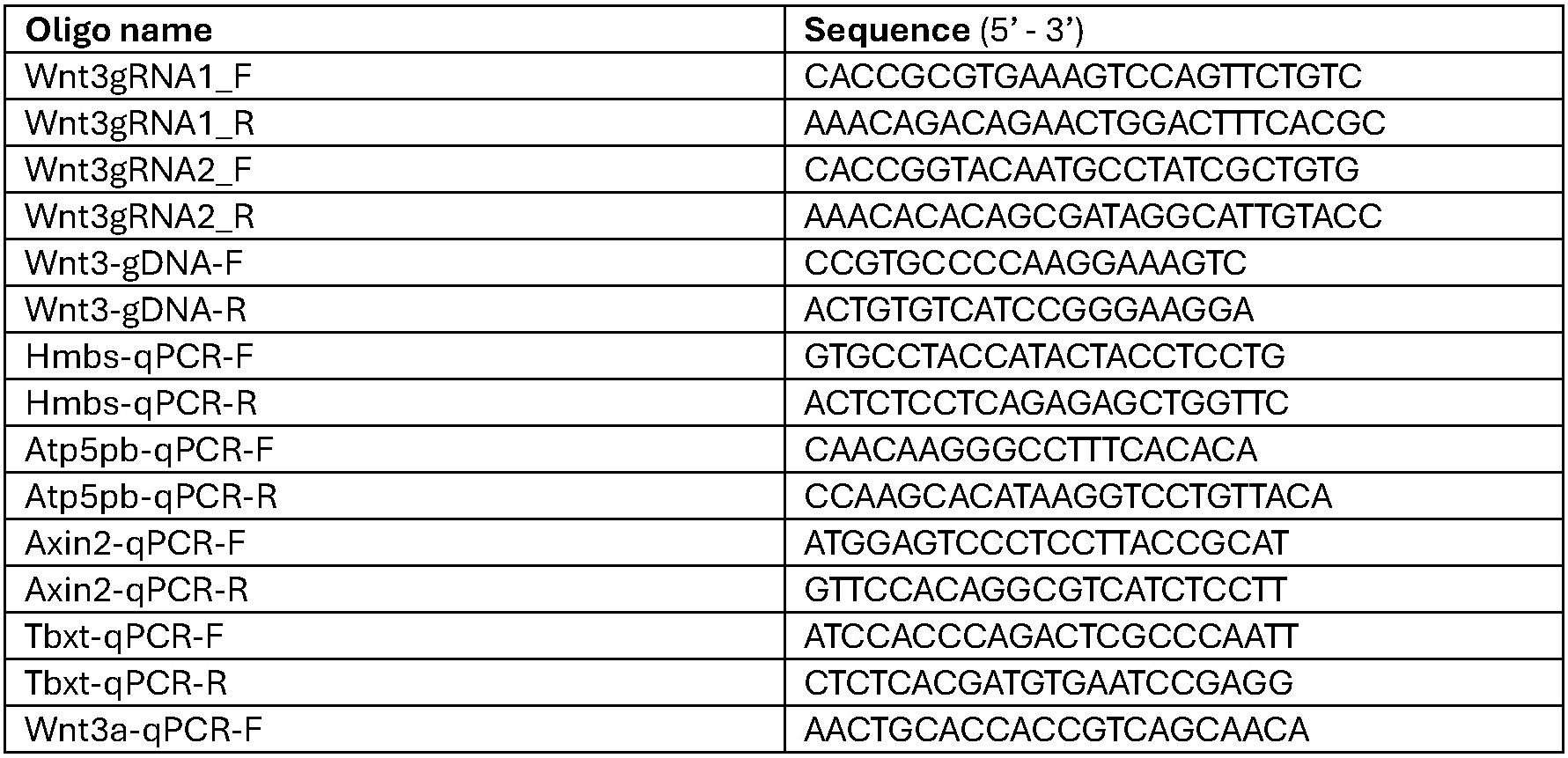
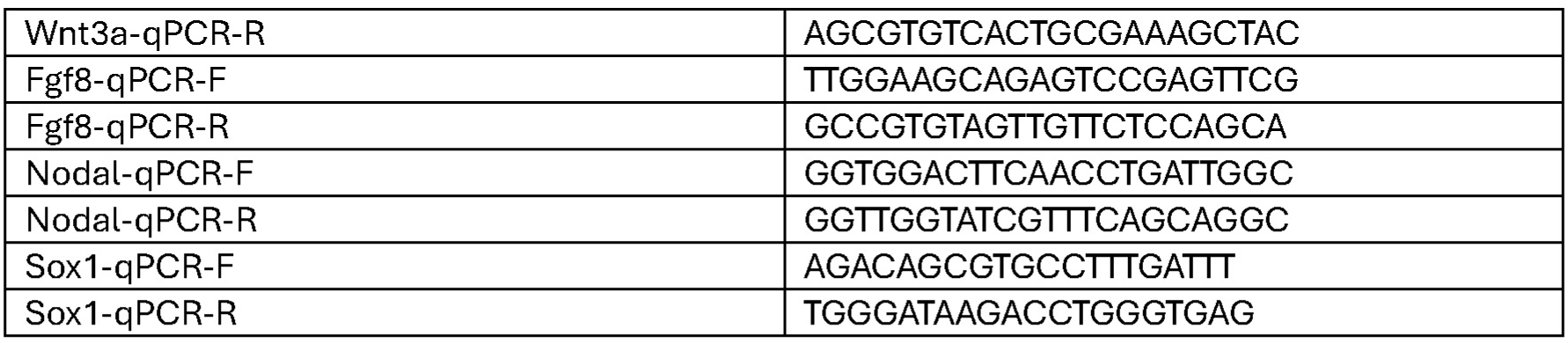
List of oligos.

**Supplementary Data Table 2.** List of antibodies.

| Primary antibodies | concentration |
| --- | --- |
| Goat anti-Otx2 (R&D Systems #AF1979) | 1:200 |
| Rat anti-Nanog (eBioscience #14-5761-80) | 1:250 |
| Rabbit anti-Cdx2 (Invitrogen #EPR2764Y) | 1:200 |
| Goat anti-Brachyruy (R&D Systems #AF2085) | 1:100 |
| Rabbit anti-Sox2 (Abcam #AB92494) | 1:200 |
| Mouse anti-Oct3/4 (BD Biosciences #611203) | 1:200 |
| **Secondary antibodies** |  |
| Donkey anti-Rabbit (A647 or A555) | 1:500 |
| Donkey anti-Mouse (A555) | 1:500 |
| Donkey anti-Goat (A488) | 1:500 |
| Donkey anti-Rat (A647) | 1:500 |

**Supplementary Data Table 3.**
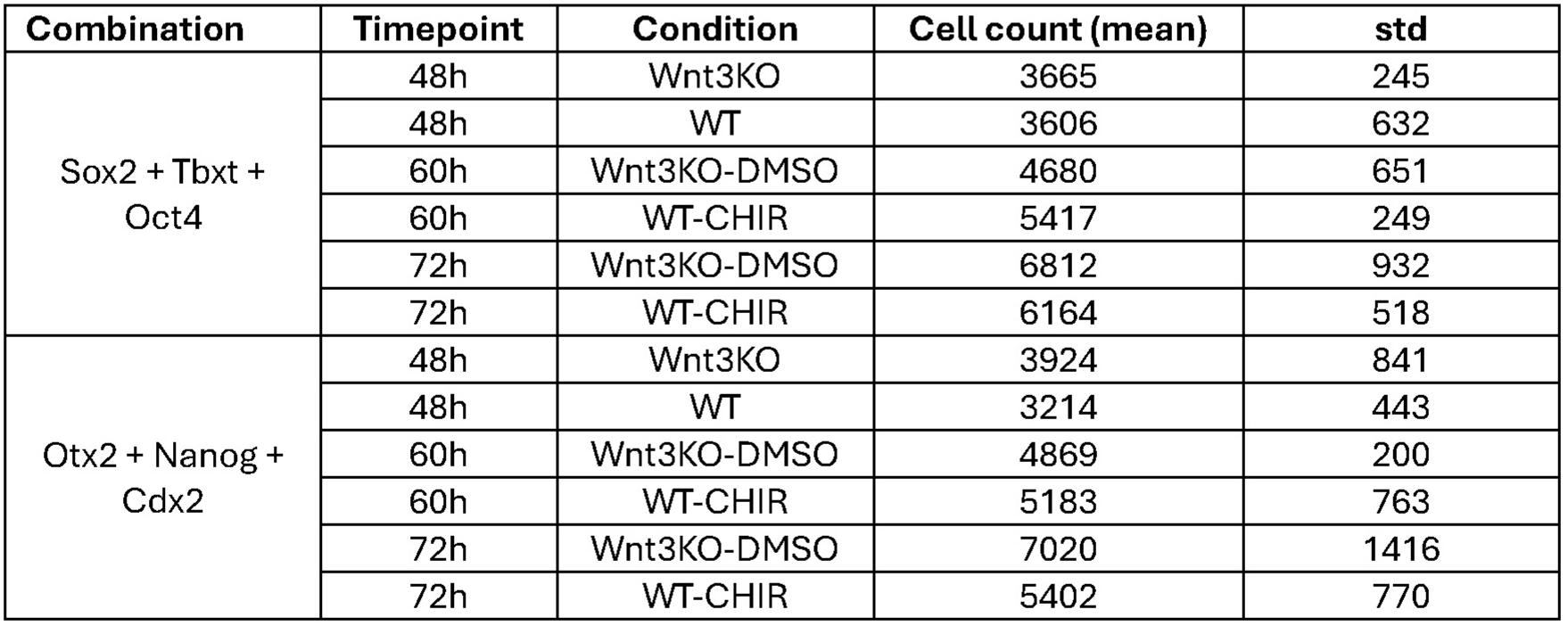
Number of 3D segmented cells per gastruloid condition/timepoint.

